# Modulation of pre-mRNA structure by hnRNP proteins regulates alternative splicing of *MALT1*

**DOI:** 10.1101/2021.10.12.464102

**Authors:** Alisha N Jones, Carina Graß, Isabel Meininger, Arie Geerlof, Melina Klostermann, Kathi Zarnack, Daniel Krappmann, Michael Sattler

## Abstract

Alternative splicing is controlled by differential binding of *trans*-acting RNA binding proteins (RBPs) to *cis*-regulatory elements in intronic and exonic pre-mRNA regions ^1-3^. How secondary structure in the pre-mRNA transcripts affects recognition by RBPs and determines alternative exon usage is poorly understood. The MALT1 paracaspase is a key component of signaling pathways that mediate innate and adaptive immune responses ^4^. Alternative splicing of *MALT1* exon7 is critical for controlling optimal T cell activation ^5,6^. Here, we demonstrate that processing of the *MALT1* pre-mRNA depends on RNA structural elements that shield the 5’ and 3’ splice sites of the alternatively spliced exon7. By combining biochemical analyses with chemical probing and NMR we show that the RBPs hnRNP U and hnRNP L bind competitively and with comparable affinities to identical stem-loop RNA structures flanking the 5’ and 3’ splice sites of *MALT1* exon7. While hnRNP U stabilizes RNA stem-loop conformations that maintain exon7 skipping, hnRNP L unwinds these RNA elements to facilitate recruitment of the essential splicing factor U2AF2 to promote exon7 inclusion. Our data represent a paradigm for the control of splice site selection by differential RBP binding and modulation of pre-mRNA structure.

---

Alternative splicing greatly expands the proteome and is associated with unique functions in metazoan organisms ^2^. Regulation of alternative splicing occurs through *cis*-acting pre-mRNA sequences, such as exonic and intronic silencers and enhancers, with cognate *trans*-acting RNA binding proteins (RBPs) ^1,3^. Pre-mRNA structure has been suggested to modulate the processing and function of RNA transcripts ^7-15^. Previous studies have shown that splicing regulation can occur through the sequestration of *cis*-regulatory RNA motifs, which upon base pairing in stem-loop structures are inaccessible to the spliceosome or cognate *trans*-acting RBPs ^14,16^. In turn, pre-mRNA structures, adopted either co-or post-transcriptionally, can be modulated by protein binding and thereby influence the accessibility of splice sites ^17,18^. However, molecular mechanisms how pre-mRNA structural elements can be modulated by RBPs to tune the level of exon inclusion or exclusion are poorly understood.

Here, we show that alternative splicing of the pre-mRNA of the mucosa-associated lymphoid tissue protein 1 (MALT1) paracaspase is regulated by an unexpected interplay of RNA structure and the RBPs hnRNP U and hnRNP L. MALT1 plays a key role in the cellular signaling pathways that promote innate and adaptive immune activation ^4,19^. The *MALT1* pre-mRNA was recently shown to express two isoforms, *MALT1A* and *MALT1B*, which only differ in the inclusion and exclusion of the 33-nucleotide long exon7, respectively (**Fig. 1a**). Notably, the inclusion of exon7 in *MALT1A* is critical for controlling T cell activation ^5^. T cell receptor engagement induces alternative splicing with inclusion of exon7 and an increase of expression of the MALT1A protein isoform in activated CD4+ T cells. Since exon7 in *MALT1A* encodes an additional TRAF6 bindingsite, enhanced TRAF6 recruitment in MALT1A ultimately augments optimal T cell activation ^5^ (**Fig. 1a**). A hypomorphic patient mutation that selectively inactivates MALT1B causes a severe immune disorder, revealing the importance of both MALT1 isoforms for human immunity ^6^. Here, we unravel the molecular mechanisms that underlie the opposing roles of two RBPs, namely hnRNP U and hnRNP L, in *MALT1* pre-mRNA splicing. Our findings serve as a paradigm demonstrating the role of RNA binding proteins in modulating pre-mRNA structure for controlling alternative splicing regulation.

**Figure 1.**
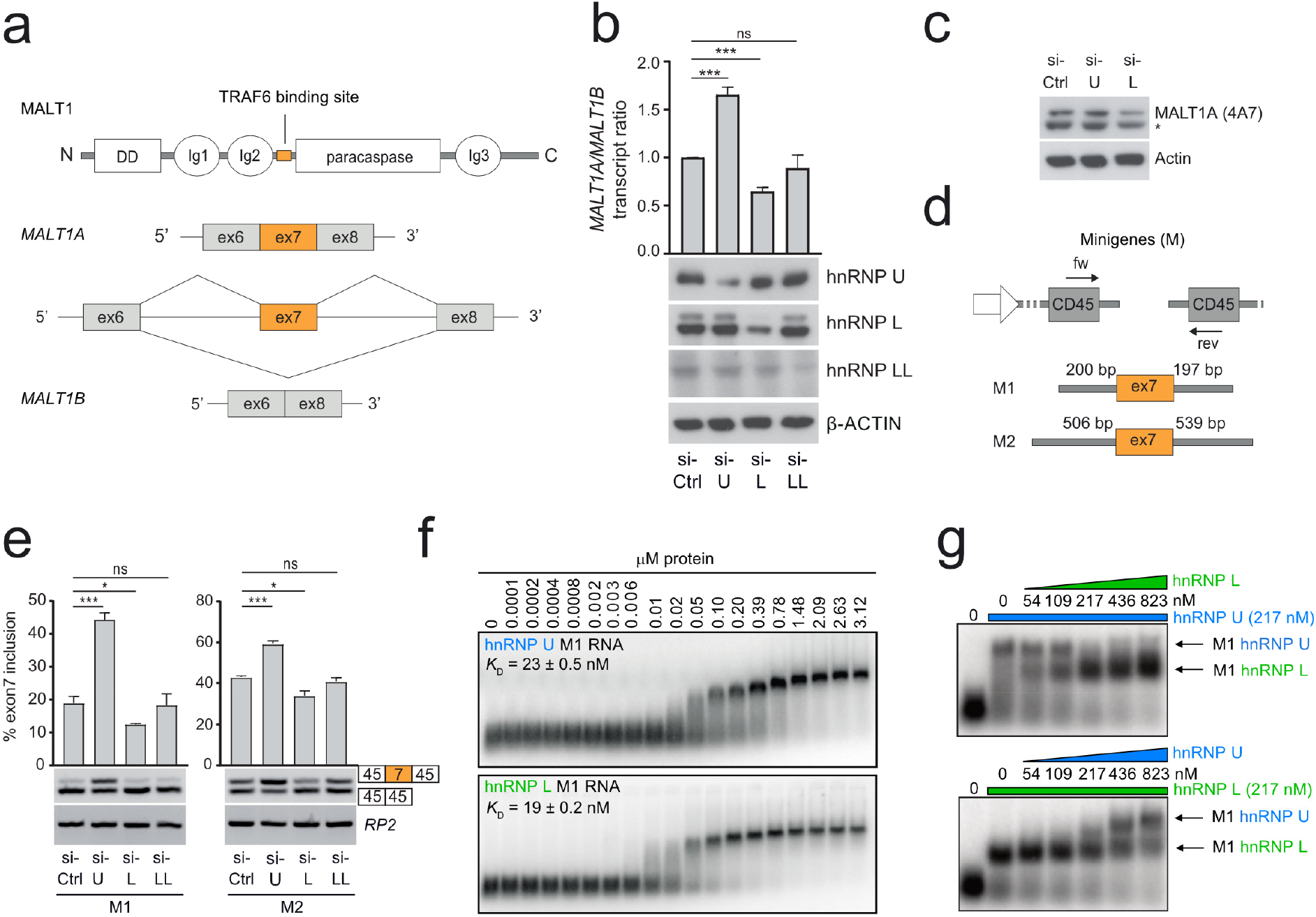
Identification of RNA elements and RBP regions regulating *MALT1* exon7 splicing. **a**, MALT1 A and B protein isoforms differ depending on inclusion or exclusion of exon7, which encodes for an 11-amino acid TRAF6 binding domain that regulates downstream function. **b**, Quantification of endogenous *MALT1* transcripts upon knockdown of hnRNP U, hnRNP L, and hnRNP LL. **c**, Endogenous protein levels of MALT1A upon knockdown of hnRNP U and hnRNP L. Asterisk indicates an unspecific band. **d**, MALT1 minigene constructs that recapitulate splicing regulation of endogenous *MALT1*. **e**, Quantification of *MALT1* splicing on minigene constructs upon knockdown of hnRNP U, hnRNP L and hnRNP LL. **f**, EMSA showing that hnRNP U and hnRNP L bind with low nanomolar affinity to the *MALT1* minigene RNA. **g**, EMSAs showing that hnRNP L and hnRNP U compete for binding to the *MALT1* minigene RNA. Data are representative for four (**b**) or three (**e**) independent experiments. Depicted is the mean ± s.d. (**b**; n = 4) or (**e**; n = 3). *p<0.05; **p<0.01; ***p<0.001; ns, not significant; unpaired Student’s *t*-test.

## Antagonistic effects of hnRNP U and hnRNP L on MALT1 splicing

Knockdown of hnRNP U in Jurkat T cells enhanced *MALT1A* transcript and protein levels, while downregulation of hnRNP L decreased *MALT1A* mRNA and protein (**Fig. 1b,c**). Note that compared to other cell lines (HeLa, U2OS and HEK293), Jurkat T cells express very low amounts MALT1A transcript and protein and hnRNP L acts as a positive regulator of exon7 inclusion and MALT1A expression in all cell lines tested (**Extended Data Fig. 1a,b**). In Jurkat T cells, the antagonistic roles of hnRNP U and hnRNP L on *MALT1* alternative splicing are recapitulated with minigenes that include ∼200 (M1) or ∼500 (M2) additional nucleotides (nt) both 5’ and 3’ flanking exon7 (**Fig. 1d,e**). Notably, deletion of intronic regions reveals that 200 nt flanking exon7 in the M1 minigene are necessary and sufficient to confer hnRNP splice factor responsiveness (**Extended Data Fig. 1c,d**). Although hnRNP L and LL are believed to serve redundant functions ^20-24^, the knockdown of hnRNP LL did not affect *MALT1* exon7 inclusion in Jurkat or other cells (**Fig. 1b, Extended Data Fig. 1b**). We conclude that hnRNP U suppresses, while hnRNP L enhances *MALT1* exon7 inclusion.

Using electrophoretic mobility shift assays (EMSAs) we found that hnRNP U and hnRNP L directly interact with the M1 pre-mRNA with similar nanomolar affinities, corresponding to dissociation constants (*K*_D_) of 23.0 ± 0.5 and 19.0 ± 0.2 nM, respectively (**Fig. 1f**). The Hill coefficient for binding of both proteins is approximately 5, indicating the presence of several binding sites for each protein (**Extended Data Table 1**). Shortening the RNA to a 200 nt fragment (M1 nt 1 - 200) covering sequences 5’ of exon7 or 231 nt (M1 nt 200 - 430) including exon7 and 3’ sequences reveals slightly weaker, yet still nanomolar binding affinity (∼30 nM) and Hill coefficients of ∼3 for both hnRNP U and hnRNP L (**Extended Data Fig. 1e, Extended Data Table 1**). Because of their antagonistic roles in *MALT1* splicing, we speculated that both RBPs do not associate simultaneously to the exon7-containing pre-mRNA. Indeed, competition EMSA demonstrated that preformed hnRNP U-M1 RNA complexes or the slightly faster migrating hnRNP L-M1 RNA complexes were displaced by increasing concentrations of free hnRNP L or hnRNP U, respectively (**Fig. 1g**). The exchange in RBP-RNA complexes occurred at approximately 1:1 stoichiometry of both RBPs, which is in line with the comparable affinities of hnRNP U and L for the M1 pre-mRNA. The absence of super-shifted RBP-RNA complexes indicates that hnRNP U and hnRNP L bind to the RNA in a mutually exclusive manner. The displacement suggests the RBPs may compete for binding to similar regions in the M1 *MALT1* pre-mRNA.

### RNA secondary structure determines MALT1 pre-mRNA splicing

We next investigated the presence of secondary structure potentially involving the exon7 splice sites in the *MALT1* pre-mRNA. Using SHAPE (selective 2′-hydroxyl acylation analyzed by primer extension) chemical probing ^25^, we show that the M1 pre-mRNA is well-structured and can be broadly split into two domains (**Fig. 2a**). Domain 1 comprises the first ∼150 nt and consists of three stem-loop (SL) RNA structures (SL0-SL3). Domain 2, which comprises the remaining ∼250 nt, consists of four stem-loops (SL4-SL8). Interestingly, SL4 harbors the poly-pyrimidine tract (Py-tract) of the 3’ splice site of the preceding intron, and the first 11 nt of exon7. Exon7 then extends into a hammerhead-like RNA structure, comprised of SL5 and SL6. SL5 harbors the 5’ splice site of exon7. The primary sequence spanning SL4 – SL6 is highly conserved across divergent species (**Extended Data Fig. 2**). The sequestration of the Py-tract and 5’ splice site flanking exon7 in these structured elements suggests that RNA structure may be directly involved in splicing regulation.

**Figure 2.**
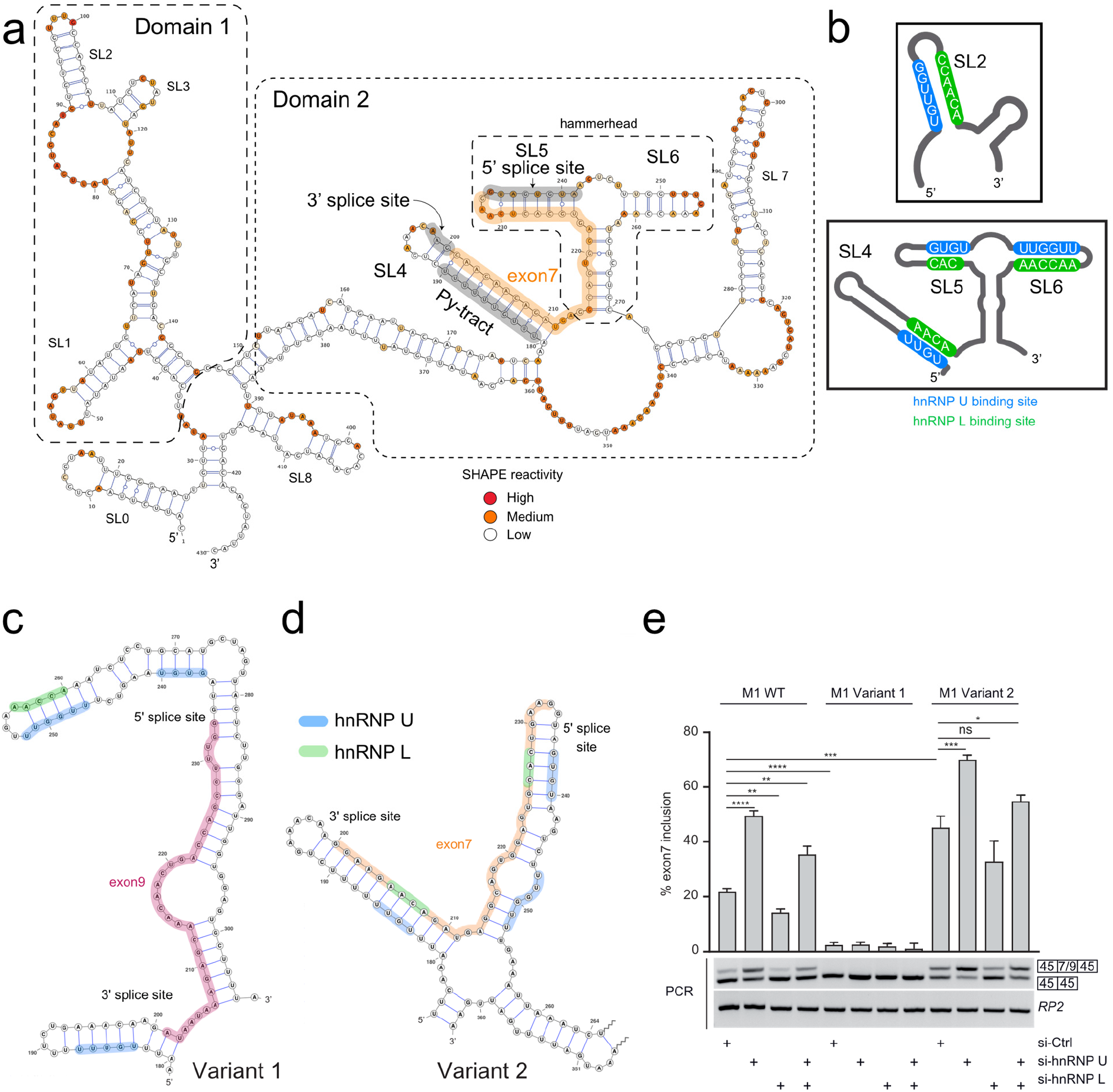
Identification of *cis*-regulatory motifs and RBP binding sites regulating *MALT1* splicing. **a**, SHAPE-derived secondary structure of the *MALT1* minigene RNA. Domains 1 and 2 are outlined, with stem-loops (SL) and splice signals highlighted and annotated. Non-reactive, semi-reactive and highly reactive nucleotides are colored white, orange, and red, respectively. **b**, Binding sites for hnRNP U (blue) and hnRNP L (green) across the *MALT1* minigene RNA. **c,d**, SHAPE-derived secondary structure of variant 1 and variant 2 of *MALT1* minigene RNAs, zoomed in to the region that harbors the 5’ and 3’ splice signals flanking exon7. **e**, Effects and quantification of splicing regulation of exon7 or exon9 (variant 1) upon single or combined knockdown of hnRNP U and hnRNP L comparing the wildtype (WT) M1, variant 1 and variant 2 minigenes. Data are representative for three independent experiments. Depicted is the mean ± s.d. *p<0.05; **p<0.01; ***p<0.001; ****p<0.0001; ns, not significant; unpaired Student’s *t*-test.

Close inspection of the primary sequences of SL2, SL4, SL5, and SL6 revealed a striking feature: All four stem-loop structures harbor complementary GU- and CA-containing RNA sequences in the two strands that base pair in the RNA helical stem (**Fig. 2b**). GU- and CA-rich sequences have been suggested to be recognized by hnRNP U and hnRNP L, respectively ^24,26,27^. Interestingly, regions in exon7 that base pair with GU-rich sequence around the Py-tract and the 5’ splice site are comprised of CA-containing sequences. Considering that these regions harbor hnRNP L and hnRNP U binding motifs, our data suggest that RNA binding by these RBPs may be directly involved in controlling spliceosome accessibility.

To investigate the functional importance of RNA primary sequence and secondary structure for exon7 splicing, we designed two variants that selectively disrupt the structure of the M1 pre-mRNA, without affecting essential splice signals (**Extended Data Fig. 3a**). In variant 1, exon7 is replaced with exon9 of *MALT1*, which has an identical length of 33 nt. SHAPE probing reveals that the flanking Py-tract and 5’ splice site sequences are sequestered in secondary structures (**Fig. 2c, Extended Data Fig. 3b**) and are thus not accessible for spliceosome assembly. Consistent with this, minigene splicing assays show that the closed conformation of *MALT1* pre-mRNA variant 1 completely prevents inclusion of exon9 (**Fig. 2e**). We rationalize that swapping to exon9 removed the exon7-encoded hnRNP L binding motifs that base pair with the Py-tract and the former SL5 in the primary variant 1 transcript (**Fig. 2c, Extended Data Fig. 3a,b**). Consistent with this, hnRNP L binding to the variant 1 pre-mRNA is slightly decreased, while hnRNP U binding remains the same compared to M1 wildtype pre-mRNA (**Extended Data Fig. 3d**). We also observed loss of splicing control by downregulation of hnRNP U or hnRNP L (**Fig. 2e**). Thus, the sequestration of the Py-tract and 5’ splice sites by exon9, in combination with the absence of hnRNP L binding motifs, renders the region inaccessible for the splicing machinery. In variant 2, we altered two nucleotides in the SL6 stem, which resulted in the destruction of SL6 and thus loss of the SL5/SL6 hammerhead structure, while maintaining the binding of hnRNP U and L (**Fig. 2d, Extended Data Fig. 3a,c,d**). Despite extended base pairing of the former SL6 with exon7 and more distant regions in the pre-mRNA, the sequestration of the Py-tract in SL4 and of the 5’ splice site in SL5 and the presence of hnRNP U and L binding motifs are retained. Minigene splicing assays with variant 2 demonstrated significantly enhanced inclusion of exon7 compared to M1 wildtype pre-mRNA (**Fig. 2e**). Importantly, the *MALT1* variant 2 is still sensitive to RBP regulation just like the *MALT1* wildtype pre-mRNA, providing evidence that SL4 and SL5, which shield the 3’ and 5’ splice sites and bind to hnRNP U and L, are critical for regulating alternative exon7 splicing.

### hnRNP L unwinds while hnRNP U stabilizes RNA structure

To investigate the mechanism by which the RBPs hnRNP U and hnRNP L regulate splicing, we performed SHAPE on the M1 pre-mRNA in the presence of each protein. Analysis of SHAPE data reveals multiple regions of differential reactivity relative to the RNA-only control indicative of specific binding regions. Upon addition of hnRNP U or L, varying degrees of differential SHAPE reactivity were observed for each of the SLs. The most pronounced effects were observed for SL5, which harbors the 5’ splice site of exon7 (**Fig. 3a**). In the presence of hnRNP L, there is an increase in SHAPE reactivity of nucleotides at the 5’ splice site. As the putative binding site for hnRNP L is opposite the 5’ splice site in the SL5 stem region, this suggests that binding by hnRNP L unwinds the stem-loop, rendering the 5’ splice site accessible. In contrast, the presence of hnRNP U results in a reduction of SHAPE-reactive nucleotides in SL5, suggesting that hnRNP U binds and stabilizes the structured elements preventing acylation by the SHAPE reagent. To support these conclusions, we used fluorescence quenching assays to monitor RNA unwinding ^28^. A FAM-fluorophore and a DABCYL quencher were conjugated to the 5’ and 3’ ends of each *MALT1* pre-mRNA SL4 and SL5, to allow detection of stem-loop opening; unwinding of the RNA reduces the quenching effect, resulting in an increase in FAM-fluorescence emission. Indeed, we observed a concentration-dependent enhancement of fluorescence emission of SL4 and SL5 by increasing the hnRNP L protein concentration, consistent with an unwinding of the RNA stem-loops (**Fig. 3b**). In contrast, no increase in fluorescence emission was observed upon binding of hnRNP U to SL4. The small fluorescence emission for SL5 at low hnRNP U amounts was not further increased by higher concentrations, showing that hnRNP U was not able to efficiently unwind both stem-loops (**Fig. 3b**). Thus, while hnRNP L is able to destabilize and unwind the regulatory *MALT1* exon7 stem-loop structures, hnRNP U is unable to do so and rather stabilizes the RNA hairpins.

**Figure 3.**
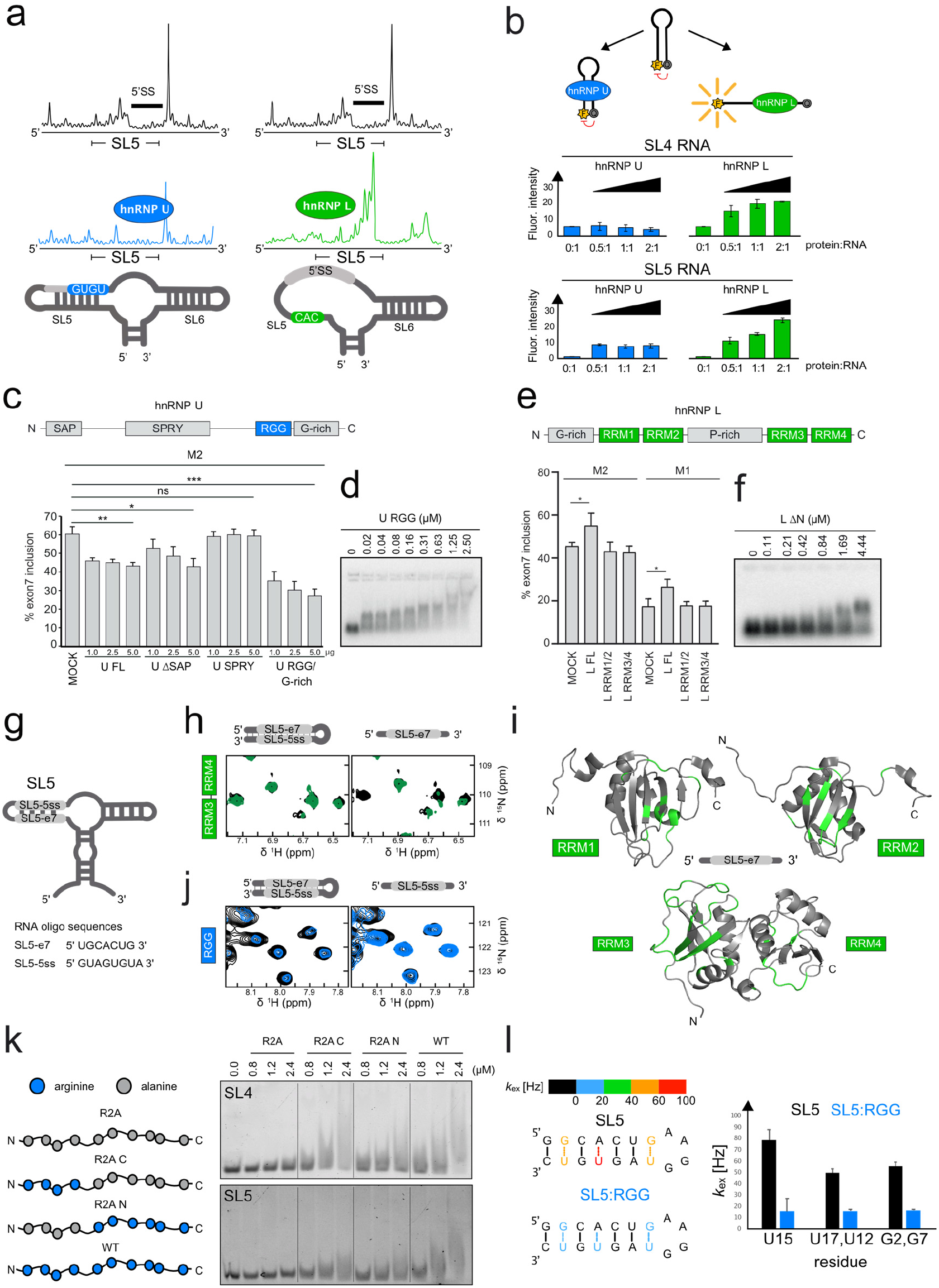
pre-mRNA structure and effect of RNA variations on exon7 splicing. **a**, Raw SHAPE reactivity traces corresponding to SL5, which harbors the 5’ splice site, in the absence of protein (black), in the presence of hnRNP L (green) and hnRNP U (blue). **b**, Fluorescence quenching assays with the SL4 RNA hairpin labeled with a fluorescent dye and quencher at the 5’ and 3’ termini show that hnRNP L unwinds, whereas hnRNP U maintains secondary structures of splice signal-containing stem-loops. Errors refer to 3 biological replicates. **c**, Minigene splicing assay quantification upon overexpression (1, 2.5, and 5 µg) of various hnRNP U constructs. **d**, EMSA of with hnRNP U RGG domain with *MALT1* M1 minigene RNA **e**, Minigene splicing assay quantification upon overexpression (5 µg) of various hnRNP L constructs. Data are representative for three (**c,e**) independent experiments. Depicted is the mean ± s.d. (**c,e**; n = 3). *p<0.05; **p<0.01; ***p<0.001; ns, not significant; unpaired Student’s *t*-test. **f**, EMSA of hnRNP L RRM1-4 with *MALT1* M1 minigene RNA. **g**, Schematic of stem-loop RNA structures and primary sequences harboring the 5’ and 3’ splice sites flanking the *MALT1* exon7. **h**, ^1^H ^15^N HSQC NMR spectra of ^15^N-labeled hnRNP L RRMs 3,4 free (black) and in the presence of equimolar concentrations of SL5 and SL5-e7 RNA (green). **i**, CSPs induced by the SL5-e7 RNA are mapped in green on the structure of hnRNP L RRM 1, RRM 2 and RRRMs 3,4; amino acids are labeled for reference (structures adapted from PDB IDs 2MQO, 2MQP, and 2MQQ). **j**, ^1^H ^15^N HSQC NMR spectra of ^15^N-labeled hnRNP U RGG free (black) and in the presence of equimolar concentrations of SL5 and SL5-e7 RNA (blue). **k**, Schematic representation of wildtype and arginine to alanine RGG mutants and corresponding binding shift assays with SL4 and SL5. **l**, Imino proton-exchange rates (*k*_ex_) from CLEANEX-PM NMR experiments for SL5 in the absence and presence of the RGG domain of hnRNP U. Error bars indicate the fitting error.

To enable biophysical and structural analyses of RBP binding to *MALT1* pre-mRNA elements, we determined the regions in the hnRNP U and L proteins necessary for controlling splicing and RNA binding. Overexpression of full-length (FL) hnRNP U decreases exon7 inclusion (**Fig. 3c**). There is a small contribution of the N-terminal SAP domain, a putative DNA binding fold, which also binds to the M1 pre-mRNA (**Fig. 3c, Extended Data Fig. 4a-d**). However, the C-terminal RNA-binding RGG/G-rich region is sufficient to strongly reduce exon7 inclusion (**Fig. 3c**). In line with this, the RGG/G-rich region and RGG domain alone bind with nanomolar affinity to the M1 pre-mRNA (**Fig. 3d, Extended Data Fig. 4d**), demonstrating that the RGG/G-rich region of hnRNP U is sufficient to confer RNA binding and repression of *MALT1* exon7 splicing. While full-length hnRNP L increases exon7 inclusion, no discernable difference in exon7 splicing is observed upon overexpression of the tandem RNA recognition motif (RRM) domains RRM1,2 or RRM3,4,indicating that all four RRMs are required for regulation of splicing in cells (**Fig. 3e, Extended Data Fig. 4e,f**). hnRNP L RRM1,2 alone does not bind strongly to the M1 pre-mRNA in EMSA, but a fragment containing all four RRMs (L ΔN) or RRM3,4 bind more readily to the M1 pre-mRNA, even though with lower affinity than full-length hnRNP L (**Fig. 3f, Extended Data Fig. 4g**). Thus, hnRNP L RRM3,4 facilitates M1 RNA binding, but all RRMs are required to promote *MALT1* exon7 inclusion.

We used NMR to characterize the binding of the SL4 and SL5 hairpins harboring the splice sites flanking exon7 and their single-stranded constituents to the tandem RRM domains of hnRNP L (RRMs 1,2 and 3,4) and the hnRNP U RGG domain (**Fig. 3g-i, Extended Data Fig. 5**). We first analyzed the interactions with hnRNP L. Relative to the structured stem-loops (SL4 and SL5), titration of single-stranded RNAs (SL4-e7 and SL5-e7) resulted in significantly stronger chemical shift perturbations (CSPs) and severe line broadening for amide resonances of hnRNP L RRM 1,2 (**Extended Data Fig. 5a,c**). Interestingly, titration of both SL4-e7 and SL4, in addition to SL5-e7, resulted in notable CSPs for amide resonances of hnRNP L RRM 3,4 (**Extended Data Fig. 5b,d**). Given that hnRNP L is known to bind to single-stranded CA motifs, this requires unwinding of SL4 (**Fig. 3b**) and is likely enabled by the reduced stability of SL4. Consistent with this SL4 harbors weak and dynamic UA base pairs, as indicated by the increased imino proton exchange detected by CLEANEX-PM ^29^NMR experiments (**Extended Data Fig. 6a,b**). Spectral changes induced by the SL5-e7 RNA binding are mapped onto the hnRNP L RRM 1, RRM 2, and tandem RRM 3,4 structures (**Fig. 3i**), and are in good agreement with the RNA binding interface previously analyzed ^30^. Notably, more significant spectral differences occur for hnRNP L RRM 3,4, which is consistent with the increased binding we observed for this construct with the M1 minigene pre-mRNA (**Extended Data Fig. 4g**). Altogether, we find that hnRNP L tandem RRMs bind to single-stranded CA-containing sequences, and are able to recognize and bind CA-containing sequences sequestered in weak and dynamic RNA secondary structure. As revealed by the fluorescence quenching assays, association of hnRNP L with SL4 and SL5 RNAs involves unfolding of the stem-loop structures.

We next characterized the *MALT1* RNA recognition by hnRNP U. Inspection of the ^1^H,^15^N NMR correlation spectra of the RGG domain of hnRNP U reveals that it is intrinsically disordered, as indicated by poor spectral dispersion of the amide resonances (**Extended Data Fig. 7a**). Titration of the single-stranded RNA motifs from SL4-PYT or SL5-5ss to ^15^N-labeled RGG domain shows only minor spectral changes and limited line broadening of amide resonances, indicating very weak, non-specific interaction (**Fig. 3j, Extended Data Fig. 7a**). Interestingly, SL5-5ss RNA induced more line broadening than SL4-PYT. Using EMSA assays, we find that the SL5-5ss is capable of forming a duplex, thus providing an artificial duplex bound by the RGG domain (**Extended Data Fig. 7a**, gel inset). In contrast, addition of the SL4 or SL5 hairpin RNAs triggers severe line broadening. While imino NMR spectra of SL4 upon addition of the RGG domain shows severe line broadening of all imino residues at a 1:1 ratio, addition of the RGG domain of hnRNP U to SL5 shows distinct chemical shift perturbations of G and U imino protons. The fact that the imino signals remain observable, although with significant line broadening, allows us to conclude that the RGG domain does not unwind RNA stem-loops (**Extended Data Fig. 7b**). Our data indicate a strong preference of the RGG domain for the structured RNAs consistent with previous reports ^26^. Using isothermal titration calorimetry (ITC), we determined that SL4 and SL5 bind to the RGG domain of hnRNP U with a dissociation constant *K*_D_ of 0.64 and 1 μM, respectively, but no association is detectable with the single-stranded sequences derived from the two stem-loops (**Extended Data Fig. 7c, Extended Data Table 2**). Considering the intrinsically disordered conformation of the RGG domain, we suspected that the binding mechanism of the RGG domain to a structured RNA may rely on electrostatic interactions of arginine side chains with the phosphate backbone of the RNA. Indeed, the interaction is strongly reduced in the presence of increasing sodium chloride concentrations, as seen by the reappearance of narrow NMR signals for both SL4 and SL5 in both amide and arginine side chain Hε resonances, indicative of RGG protein release from the RNA (**Extended Data Fig. 7d,e**). Clearly, the RGG domain of hnRNP U does not efficiently bind to GU-rich sequence motifs in single-stranded RNA, but recognizes GU sequences in the context of a double-stranded RNA, demonstrating that hnRNP U acts by binding and stabilizing RNA structural elements.

Having established that the RGG domain of hnRNP U binds to structured RNA stem-loops, we carried out binding shift assays of the RGG domain with SL4 and SL5, mutating either all, the four N-terminal, or the five C-terminal arginine residues to alanines (**Fig. 3k**). We found that binding is significantly reduced when only half of the arginines are present, and completely abolished with the arginine to alanine mutation of all arginine residues (**Fig. 3k**). Thus, arginines are responsible for facilitating binding of the RGG domain with RNA stem-loop elements.

We further validated the RGG domain’s stabilizing effect on structured RNA elements by monitoring RNA imino proton exchange in the absence and presence of the RGG domain using CLEANEX-PM NMR experiments (**Fig. 3l, Extended Fig. 8a,b**). Here, in the absence of the RGG domain, imino signals in SL5 (i.e. of G2, G7, U12, U15, and U17) rapidly exchange with water, with *k*_ex_ rates as high as 80 Hz. However, the rates of exchange in these corresponding imino protons decrease by nearly 4-fold in the presence of the RGG domain of hnRNP U. These results indicate a limited stability of the SL5 hairpin and support the notion that the RGG domain stabilizes the RNA secondary structure upon binding.

### Unwinding of RNA structure by hnRNP L promotes recruitment of U2AF2

The essential splicing factor U2AF2 is required for the recognition of the Py-tract motif in the 3’ splice site of pre-mRNA introns. Biochemical and structural studies have shown that the RRM1,2 tandem domains in U2AF2 are necessary and sufficient to recognize single-stranded Py-tract RNAs ^31,32^. However, in the context of the *MALT1* pre-mRNA, the Py-tract of exon7 is sequestered in the secondary structure of SL4, posing the question of how U2AF2 can access this region. In fact, U2AF2 RRM1,2 does not bind full-length M1 *MALT1* pre-mRNA or the structured SL4 RNA at protein concentrations up to 6 µM (**Fig. 4a**, top**; Extended Data Fig. 9a**). In contrast, EMSAs with U2AF2 RRM1,2 and a single-stranded Py-tract show nanomolar binding (**Fig. 4a**, bottom). ITC data confirmed that U2AF2 RRM1,2 binds to the single-stranded Py-tract with a *K*_D_ of 2.7 ± 0.7 µM, while there is no detectable binding when the Py-tract is sequestered in RNA structure of the SL4 (**Fig. 4b, Extended Data Table 2**). Consistent with this, an NMR titration of ^15^N-labeled U2AF2 RRM1,2 with SL4 RNA shows negligible spectral changes, while significant line broadening observed upon addition of the single-stranded Py-tract RNA indicates a strong interaction (**Fig. 4c**).

**Figure 4.**
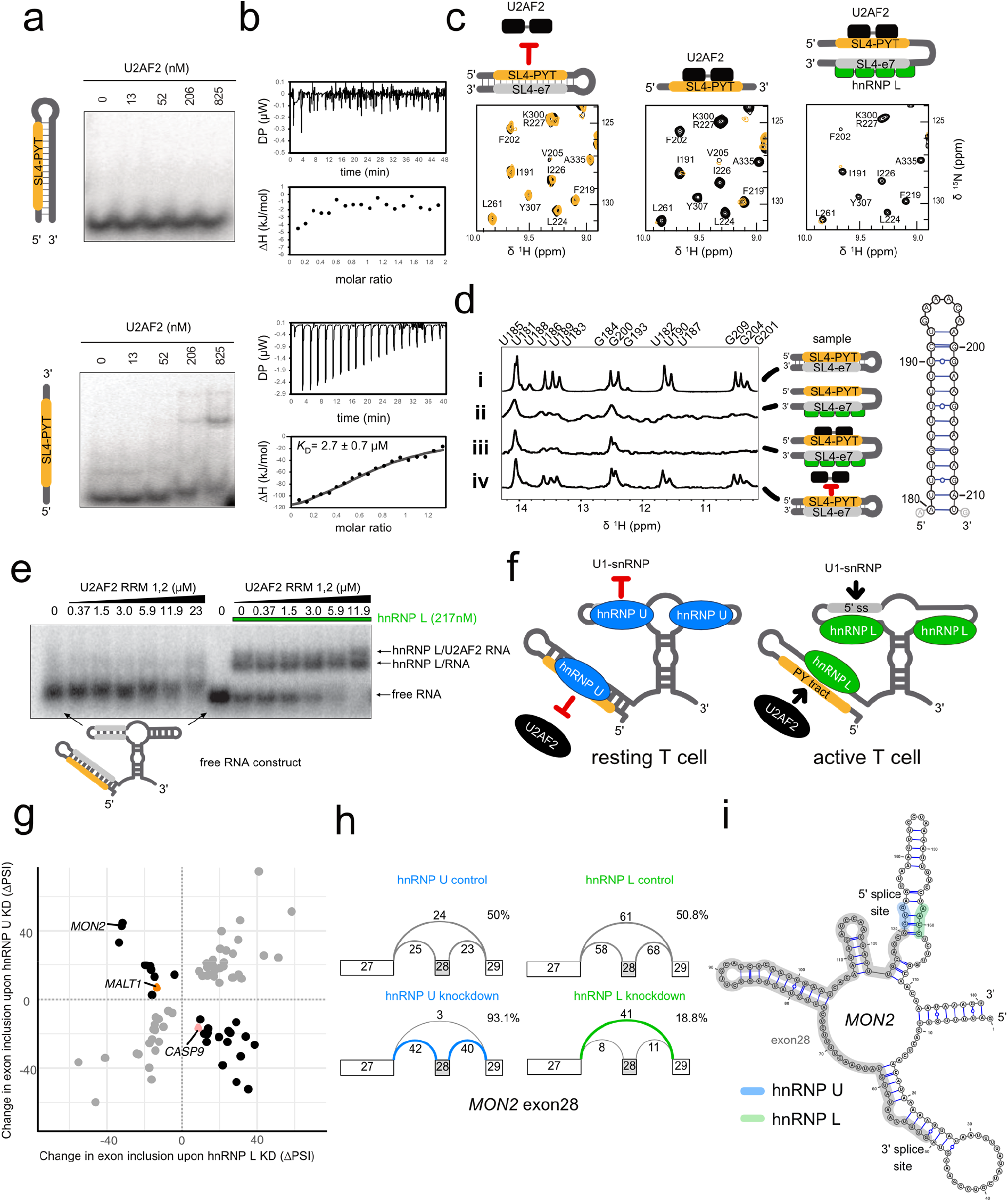
Molecular mechanisms of differential exon7 splicing regulation by hnRNP U and hnRNP L. **a**, EMSA of U2AF2 RRM1,2 with the single-stranded poly-pyrimidine (Py)-tract or SL4 sequestering the Py-tract. **b**, ITC data for binding of U2AF2 RRM1,2 with the single-stranded Py-tract and the structured SL4 RNA. **c**, ^1^H ^15^N HSQC NMR spectra of ^15^N-labeled U2AF2 RRM1,2 with SL4 RNA (left), the single-stranded Py-tract (middle), and in the presence of hnRNP L RRM1-4 and SL4 RNA (right). **d**, 1D ^1^H imino NMR spectra of SL4 RNA free (i) in the presence of hnRNP L (ii), of hnRNP L and U2AF2 RRM1,2 (iii) and of U2AF2 RRM1,2 only (iv). **e**, EMSA of U2AF2 RRM1,2 in the absence (left) and presence (right) of hnRNP L RRM1-4 with SL4-hammerhead RNA construct. **f**, Overview of the proposed mechanism of splicing regulation of *MALT1* exon7 by hnRNP U and hnRNP L in T cells. **g**, Alternative cassette exons regulated by both hnRNP U and hnRNP L in HepG2 cells. Changes in relative abundance of the exon inclusion junction are shown for all exons that are significantly regulated by both RBPs (> 5% change in percent selected index [ΔPSI] for at least one junction, probability > 90%). *MALT1* exon7 and *CASP9* exon3,4,5,6 (probability > 80%) are shown in orange and pink, respectively. **h**, Skipping of exon28 of *MON2* (RefSeq transcript NM_015026.3) and its dependence on hnRNP U and hnRNP L is shown from shRNA knockdown data available from ENCODE and processed in MAJIQ. **i**, The secondary structure of *MON2* (with exon28 traced in grey, the potential hnRNP U/5’ splice site traced in blue, and the potential hnRNP L binding sequence traced in green).

We next used NMR to determine if U2AF2 RRM1,2 binding to the Py-tract that is inaccessible in SL4 could be primed by hnRNP L. In fact, severe line broadening in the NMR spectra of U2AF2 RRM1,2 upon addition of hnRNP L to a preformed U2AF2 RRM1,2-SL4 complex (**Fig. 4c**) is consistent with Py-tract binding to U2AF2 that is enabled by the binding (and unwinding) of SL4 by the strong interaction of hnRNP L with CA-rich motifs in SL4 (SL4-e7, **Extended Data Fig. 5a,b**). To monitor the conformation of the SL4 RNA (**Fig. 4d**, right) we compared 1D NMR spectra of the imino region of SL4 in the absence and presence of U2AF2 RRM1,2 alone. The imino signals observed for the free RNA (**Fig. 4d**, i) demonstrated the presence of a folded hairpin structure. Binding by hnRNP L leads to severe line broadening of the imino signals, consistent with a at least partial unwinding of the SL4 RNA (**Fig. 4d**, ii). This is also seen in the presence of hnRNP L and U2AF2 (**Fig. 4d**, iii), where the two RBPs presumably bind to the single-stranded SL4-7 and SL4-PYT RNA regions. In contrast, imino signals of SL4 RNA are unaffected in the presence of only U2AF2 (**Fig. 4d**, iv). This shows that the RNA remains folded and that U2AF2 alone is unable to unwind the SL4 structure. Further, inspection of a 2D imino NOESY experiment showed that intra-residue nuclear Overhauser effects (NOEs) for most base pairs are no longer observed. Resonances that remain correspond to nucleotides in the upper region of the stem. This suggests that RNA base pairing in the stem region is severely disrupted, resulting in an at least partial unwinding of the SL4 (**Extended Data Fig. 9b**). The fact that the 1D imino signals of the SL4 RNA in the presence of U2AF2 RRM1,2 and hnRNP L are very similar to those in the presence of hnRNP L alone indicates that hnRNP L binding at least partially opens SL4 and thereby primes for U2AF2 binding.

To confirm these conclusions in the presence of the pre-mRNA, we performed EMSA experiments monitoring binding of hnRNP L and U2AF2 to a *MALT1* pre-mRNA comprising exon7 and the stem-loops 4-6 (SL4-hammerhead RNA). By adding increasing concentrations of U2AF2 RRM1,2 to the preformed hnRNP L SL4-hammerhead RNA, a ternary hnRNP L-RNA-U2AF2 complex is formed, showing that hnRNP L facilitates association of U2AF2 to the Py-tract (**Fig. 4e**). Similarly, we observe a slower migration of the M1 minigene RNA when the concentration of hnRNP L is held constant and the concentration of U2AF2 is increased (**Extended Data Fig. 9c**). Thus, by binding to CA-rich RNA elements in exon7 of *MALT1* pre-mRNA, hnRNP L unwinds SL4 and thereby facilitates U2AF2 to associate with the single-stranded Py-tract. This is enabled by the strong, nanomolar binding affinity of hnRNP L to single-stranded CA-rich RNA motifs (**Fig. 1f**).

Binding of hnRNP U stabilizes structured RNA motifs and thus inhibits unwinding. To support this regulation by unwinding of RNA secondary structure by a single-stranded RBP, the RNA stem regions involved (i.e. SL4 and SL5) must exhibit reduced thermodynamic stability. This is in fact reflected by the sequence composition of the *MALT1* exon7 RNA stem-loop structures. The duplex regions of these stem-loop structures are mainly formed from base-pairing of CA- and GU-rich sequences, involving many AU base pairs (**Fig. 2a,b**). The reduced thermodynamic stability enables binding of hnRNP L by capturing a minor fraction of single-stranded RNA conformations that preexists for the weakly based-paired stem regions. In light of this, the previously reported preference of hnRNP U for GU-rich sequences thus likely reflects the preference for stem-loop structures with weak base pairing, and, in fact, does not relate to a preference of hnRNP U to bind (GU-rich) single-stranded sequences.

Our results present a new paradigm for the control of alternative splicing by pre-mRNA secondary structure, which in turn is regulated by the binding of two RBPs, hnRNP U and hnRNP L, as shown here for the example of *MALT1* splicing. These two RBPs differentially modulate the accessibility of the splice sites of the *MALT1* alternative exon7. Splice signals are base-paired and thus inaccessible in the presence of hnRNP U, binding of hnRNP L unwinds the RNA, and facilitates recruitment of spliceosome factors (**Fig. 4f**).

### Further splicing events are regulated by hnRNP U and hnRNP L

To explore whether antagonistic regulation by hnRNP U and hnRNP L occurs at other exons in the transcriptome, we quantified alternative splicing events in hnRNP U and hnRNP L shRNA knockdown RNA-seq data for HepG2 cells available from the ENCODE database ^33^. Even though *MALT1* exon7 itself does not pass the significance thresholds due high variability (**Extended Data Fig. 10a**), we detect 27 other exon skipping events that are antagonistically regulated by both hnRNP U and hnRNP L (out of 78 shared exon skipping events; > 5% change in junction usage, probability > 90%; **Fig. 4g, Extended Data Fig. 10b,c**). Among these, about a third are regulated in the same direction as *MALT1* exon7, while the remainder follow an inverse pattern such that hnRNP U promotes inclusion, as has been reported for the exon3,4,5,6 cassette in *Caspase9* (*CASP9*) ^34^. Strongest net effects in the direction of *MALT1* exon7 are observed for exon28 in the *MON2* pre-mRNA (**Fig. 4h**). Notably, secondary structure prediction reveals that the 5’ splice site of exon28 in *MON2* is sequestered in secondary structure with a CCAA-containing sequence (**Fig. 4i**), suggesting that similar mechanisms underlie antagonistic regulation by hnRNP L and hnRNP U also for this splicing event.

As exemplified for alternative splicing of *MALT1* exon7, our results show how splicing regulation can be achieved by the modulation of RNA structure that sequesters essential splice signals. This has important implications for the interpretation of disease-associated mutations in clinical studies. Even though current estimates suggest that up to 50% of pathogenic single nucleotide polymorphisms (SNPs) are related to splicing ^35,36^, only a minor fraction of these splice-altering mutations can be mechanistically explained. As a consequence, clinical scoring schemes such as Alamut (Interactive Biosoftware, Rouen, France) are largely confined to known splice-regulatory sequence motifs. However, 15% of SNPs were found to alter local RNA structure ^10^. A detailed knowledge about RNA structures and their modulation by RBP binding will therefore be critical to improve our understanding of disease-associated splicing defects.

Our study suggests that modulation of pre-mRNA structure by the *trans*-acting RBPs hnRNP U and hnRNP L may serve as a mechanism that controls the accessibility of alternatively spliced exons for the basic splicing machinery. These findings offer an inverse perspective to multiple studies that investigated the impact of RNA structure on RBP binding ^37-39^and opens new avenues for a more holistic view of the dynamic interplay of RNA structure and *trans*-acting RBPs.

## Methods

### RNA preparation, transcription and purification

The DNAs encoding for the RNAs transcribed in this study were either generated in-house (subcloned by PCR from *MALT1A* using primers containing the T7 polymerase promoter region), or purchased as single stranded DNA templates (and supplemented with an equal amount of T7 promoter primer) from Eurofins Genomics (Eurofins). Transcription reactions were carried out in the presence of 600 ng (PCR) or 8 µM (Eurofins) DNA template, 40 mM MgCl_2_, 8 mM of each rATP, rUTP, rGTP, and rCTP, 20X transcription buffer (Tris-HCl pH 8, 100 mM Spermidine, 200 mM DTT), 5% PEG 8000, and 0.03 mg of T7 polymerase. The transcription reaction was incubated at 37°C for 3 hours, followed by denaturing purification on 6.5 - 20% urea polyacrylamide gels. The RNA was then excised and extracted from the gel by electroelution. The extracted RNA was equilibrated against a NaCl gradient (1 M, 0.5 M, 0.25 M, 0 M) followed by equilibration into a buffer containing 25 mM sodium phosphate pH 6.4, and 15 mM NaCl. All single stranded RNAs shorter than 10 nucleotides were purchased from Eurofins.

### Protein expression and purification

The full length and subdomains of hnRNP U/L, as well as RRMs 1,2 of U2AF2 were cloned into the pETM-11 vector yielding constructs with an N-terminal, TEV protease cleavable His_6_-tag. The proteins were expressed in either BL21 (DE3) or Rosetta 2 (DE3) *E. coli* strains and cultured in LB or ZYM 5052 auto-induction media. ^15^N labeled proteins were cultured in 1M9 minimal media. Cell were lysed by sonication in buffer A, containing 50 mM Tris-HCl, 1 M NaCl, 10 mM MgCl_2_, 10 mg/ml DNaseI, 1 mM AEBSF.HCl, 0.2% (v/v) NP-40, 1 mg/ml lysozyme, 0.01% (v/v) 1-thioglycerol, pH 8.0, and the lysate was clarified by centrifugation (48,000 x g) and polyethylenimine (PEI) was added to a final concentration of 0.5% (v/v) to remove the excess nucleotides (for full length constructs). After centrifugation, ammonium sulfate was added to the supernatant to 90% saturation to precipitate all proteins and remove the excess of PEI. Protein was then purified by immobilized metal affinity column (IMAC) purification in buffer A with an increasing imidazole gradient (50 mM to 300 mM), followed by TEV cleavage. The cleaved protein was further purified with IMAC. Proteins were then concentrated and purified using size exclusion chromatography (SEC). Following SEC, proteins prepared for binding shift assays were equilibrated in buffer containing 50 mM Tris-HCl, 300 mM NaCl, 0.01% (v/v) 1-thioglycerol, pH 8.0, whereas proteins prepared for NMR were equilibrated in buffer containing 25 mM sodium phosphate, pH 6.5, 150 mM NaCl and 5 mM DTT.

### Electrophoretic mobility shift assays

The RNAs were dephosphorylated with CIP (New England Biolabs, NEB) and rephosphorylated with γ-^32^P-ATP using T4 PNK (NEB) according to manufacturer instructions. RNA was diluted to 5 nM and mixed with varying concentrations of protein (as indicated in the Figures) in buffer containing 50 mM Tris-HCl, 300 mM NaCl, 0.01% (v/v) 1-thioglycerol, pH 8.0, prior to loading on a gel. 0.7% agarose gels prepared in TBE were run at 25°C in 1X TBE buffer for 1 hour at 60 V. The gels were dried without heat under a vacuum for 1 hour on top of nylon membrane, and then exposed to a phosphor plate for 3 hours prior to scanning using a Typhoon imager. Bands were quantified using Image J and the binding affinity and Hill coefficients were calculated in Kaleidagraph after fitting with to the expression: 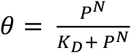, where *θ* = fraction of RNA bound, P = protein concentration, *K*_D_ = dissociation constant, and N = Hill coefficient.

### Selective 2’ hydroxyl acylation and primer extension

SHAPE, using in-house prepared 1M7 ^40^, was performed as previously described ^41^. In short, RNAs were refolded for 30 minutes at 37°C in buffer containing 100 mM NaCl, 50 mM Hepes pH 8, and 16.5 mM MgCl_2_, incubated with 5 mM 1M7 for five minutes at 37°C, followed by an ethanol precipitation; pelleted RNA was resuspended in RNase-free water. For SHAPE assays performed in the presence of protein, excess hnRNP U or hnRNP L was added to the RNA just prior to addition of 1M7. Reverse transcription was performed on the 1M7-modified RNA using a 5’-labeled 6 FAM fluorescently labeled primer (Eurofins) and the Superscript III reverse transcriptase, according to manufacturer instructions, followed by an ethanol precipitation. cDNA fragments were dissolved in HiDi formamide, followed by capillary electrophoresis analysis using an ABI 3730 Sanger Sequencer. The resulting files were analyzed with QuSHAPE to obtain SHAPE reactivities. The shotgun secondary structure (3S) method was used to validate the structure of the two independently folded domains ^42^. Transcribing domain 1 and 2 as separate transcripts, followed by SHAPE, reveals chemical probing profiles that are in agreement with those of the full-length transcript (Pearson R = 0.70 and 0.84, respectively). The Pearson R correlation coefficients were determined using the Pearson Correlation Coefficient calculator provided at (https://www.socscistatistics.com/). Secondary structure predictions, under the constraints of the SHAPE data, were carried out using RNAStructure using default folding conditions. Final RNA structural models were rendered using VARNA. SHAPE reactivity data available upon request.

### Comparative sequence analysis

We used NCBI BLASTN to curate a list of seven divergent species (very little of the *MALT1* pre-mRNA primary sequence is conserved), as shown in **Extended Data Fig. 2** ^43^. We utilized CMfinder through the webserver for aligning RNAs (WAR), which employs comparative sequence analysis to identify conserved structured motifs ^44,45^.

### NMR spectroscopy

All NMR titrations were carried out using either 100 µM unlabeled RNA (for ^1^H 1D imino experiments) or 90 µM ^15^N-labelled protein (for ^1^H,^15^N HSQC, HISQC ^46^, or SOFAST-HMQC ^47^ experiments) on 600, 800, or 900 MHz spectrometers equipped with cryogenic probes (Bruker) at 298 K.

Spectra were processed and analyzed using NMRpipe ^48^and CCPN ^49^. Resonance assignments for RRMs 1-4 of hnRNP L were completed using standard triple resonance experiments (CBCACONH, HNCACB, HNCO, and HNCACO) ^50^ and further supported by assignments reported in the BMRB for the apo RRM domains (BMRB 25038: RRM1, BMRB 25039: RRM2, and BMRB 25040: RRMs 3,4). Chemical shift perturbations were calculated as: CSP = ((Δδ^1^H)^2^+ Δδ^15^N)/6.51)^2^)^0.5^.

Imino proton resonance assignments for SL4 and SL5 RNAs were carried out using 2D ^1^H-^1^H NOESY and natural abundance ^1^H,^15^N SOFAST-HMQC experiments, recorded at 280 K and 298 K with mixing times of 80 and 120 ms.

Protein exchange rates were obtained from CLEANEX-PM experiments ^29^. Peaks were picked using TopSpin® (Bruker), signal intensities were normalized and fit using the following equation:

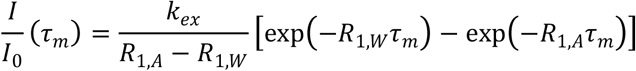

### Isothermal titration calorimetry (ITC)

Isothermal calorimetry was carried out using a MicroCal PEAQ-ITC calorimeter. Both RNA and protein were equilibrated against buffer containing 25 mM sodium phosphate, 150 mM NaCl and 5 mM DTT. RNAs were diluted to 20-30 µM and snap cooled, prior to addition to the cell. Protein (200 - 300 µM) was titrated into RNA in 2 µL increments over the course of 45 minutes at 25°C. ITC curves were fit with MicroCal PEAQ-ITC software.

### Fluorescence assays

Synthetic RNA corresponding to stem-loops SL2, SL4, SL5, and SL6 of the M1 minigene were purchased from Eurofins Genomics harboring a 5’ terminal 6 FAM fluorescent tag, and a 3’ terminal DABCYL quencher. RNAs were diluted to 400 nM, snap cooled (heated to 95°C for 5 min, followed by rapid cooling on ice for 10 min) and increasing concentrations of protein (as indicated in the figures) were added. Fluorescence emission was measured using a SpectraMax plate reader at 25°C; 6 FAM fluorescence was excited at 495 nm with a slit of 2 nm. Emission was recorded at 525 nm with a slit of 3 nm for 0.5 s (integration time).

### Cell culture and cell transfection

Jurkat T cells were cultured in RPMI 1640 medium (Life Technologies) and HEK293, HeLa and U2OS adherent cells in DMEM (Life Technologies) supplemented with 10% fetal calf serum (FCS) and 100 U/ml penicillin/streptomycin (P/S, Life Technologies) For knock-down experiments, cells were transfected with 50 – 100 nM siRNA and Atufect transfection reagent (Silence Therapeutics; Jurkat and HeLa) or Lipofectamine RNAiMAX reagent (Thermo Scientific; U2OS) and analyzed after 72 h. siRNA knockdown in HEK293 cells was performed using standard calcium phosphate transfection protocols. For minigene assays, 48 h after siRNA transfection, 2.5×10^6^ Jurkat T cells were electroporated with 2 μg minigene constructs using 220 V and 1,000 mA (Gene pulser X,Bio-Rad). After 24 h incubation, cells were lysed in protein or RNA lysis buffer. The following siRNAs were used: ON-TARGETplus Non-targeting pool (si-control), ON-TARGETplus SMARTpool si-hnRNP U, ON-TARGETplus SMARTpool si-hnRNP L, and ON-TARGETplus SMARTpool si-hnRNP LL (all from Dharmacon).

### RNA preparation, minigene assay and qPCR

RNA was isolated (InviTrap Spin Universal RNA kit, Stratec) and reverse transcribed (Verso cDNA synthesis kit, Thermo Fisher). To analyze *MALT1* exon7 inclusion or exclusion of the different minigenes, two specific vector backbone primers (*CD45* exon3 forward and *CD45* exon7 reverse) were used to amplify alternatively spliced minigene products. As internal control, RP2 levels were used. Semi-quantitative PCR was performed using Taq DNA Polymerase (NEB) and 15 ng cDNA.

To determine endogenous *MALT1A/B* levels, semi-quantitative PCR with 30 ng cDNA was performed using LongAmp® Taq DNA Polymerase (NEB) with primers in flanking exons detecting both isoforms *MALT1A* (146 bp) and *MALT1B* (113 bp).

qPCR was performed on a LightCycler 480 from Roche using LightCycler SYBR Green I Master Mix. PCR products were analyzed on 3% agarose gels. A list of all primers used for qPCR and minigene assays can be found in **Extended Data Table 3**.

### Western blot

Proteins were transferred onto PVDF-membranes for immunodetection using electrophoretic semi-dry transfer system. After transfer, membranes were blocked with 5% bovine serum albumin (BSA) or 5% milk for 1 h at room temperature and incubated with specific primary antibody (dilution 1:1,000 in 2.5% BSA/PBS-T or milk/PBS-T) overnight at 4°C. Membranes were washed in PBS-T before addition of HRP-coupled secondary antibodies (1:5,000 in 1.25% BSA or 1.25% milk in PBS-T; 1 h, room temperature). HRP was detected by enhanced chemoluminescence using the LumiGlo reagent (Cell Signaling) according to the manufacturer’s instructions. A list of all antibodies used for Western blot assays can be found in **Extended Data Table 4**.

### RNA-seq data analysis

We used MAJIQ ^51^ (version 2.2) to identify and quantify local splice variants (LSVs) in RNA-seq data from the ENCODE database. BAM alignment files (processed by STAR) of shRNA knockdown (KD) experiments for both hnRNP U and hnRNP L in the HepG2 cell line (2 replicates per condition) were retrieved from the ENCODE data portal (https://www.encodeproject.org/) via the accession numbers ENCFF764HLG, ENCFF915OWV (hnRNP L KD), ENCFF371TBZ, ENCFF403KGR (hnRNP L control), ENCFF197CGS, ENCFF451GID (hnRNP U KD), and ENCFF197UJB, ENCFF289WR (hnRNP U control). Index files were generated using Integrated Genome Browser (IGV, Broad Institute; http://software.broadinstitute.org/software/igv/). First, a MAJIQ splice graph was built on the combined BAM files from all conditions and GENCODE gene annotation (v24, human genome version hg38). The difference in junction usage (in delta percent selected index, ΔPSI) between KD and corresponding control samples was calculated with a minimum reads threshold of 3. MAJIQ VOILA was then used to calculate probabilities for each junction in the LSVs by testing for |ΔPSI| > 0.05. The VOILA output was then processed in R as follows: LSVs with at least one junction with |ΔPSI| > 0.05 and *P* value > 0.9 were considered as significant. LSVs with more than two junctions were reduced to binary events, where possible, by selecting the two main junctions with the highest positive and negative ΔPSI. For redundant LSVs that corresponded to the same splicing event from a source and target exon perspective, we kept the LSV with the highest |ΔPSI|. This procedure yielded a total of 1,719 and 1,301 significant alternative splicing events upon KD of hnRNP U and hnRNP L, respectively. We classified these LSVs as (i) exon skipping when the two main junctions connected to three exons, (ii) intron retention if reported by MAJIQ VOILA, (iii) alternative splice site when the two main junctions connected to two exons (3’ and 5’ alternative splice sites were defined via the non-overlapping junction edges), or (iv) other if they could not be unambiguously assigned. For exon skipping events, the shorter of the two junctions as assigned as inclusion junction. Fisher’s exact test for count data was performed on the overlap of significantly changing LSVs in the hnRNP U and hnRNP L KD. The LSVs corresponding to *MALT1* exon7 (LSV ID ENSG00000172175.12:s:58709976-58710072) and *CASP9* exon3-6 (LSV ID ENSG00000132906.17:s:15518110-15518395) showed the expected trend, even though did not pass the confidence threshold of *P* value > 0.9, and were re-added for the comparison of splicing changes in **Fig. 4g**.

## Acknowledgements

We would like to thank Dominik Lenhart for preparation of 1M7 SHAPE reagent and Albrecht Bindereif for hnRNP LL plasmid, Sam Asami and Gerd Gemmecker for support with NMR experiments, and the Niessing and Buchner labs for access to isotope-lab space. We are grateful to Jadiel Wasson and Mario Keller for assistance with R scripts for data analysis and Gregory Wolfe for assistance with image rendering. We thank Oliver Rossbach, Julian König, Andreas Schlundt, Florian Heyd, Juan Valcárcel, and members of the Sattler group for helpful discussions. We acknowledge access to NMR measurements at the Bavarian NMR center.

## Author contributions

ANJ performed and analyzed EMSA and ITC experiments, SHAPE chemical probing and RNA structural modeling, NMR experiments and fluorescence assays. CG and IM performed in cell assays. AG and ANJ expressed and purified protein constructs used in this study. ANJ, MK, and KZ carried out genome-wide analysis of splicing events. ANJ, DK and MS designed the study, analyzed data and wrote the manuscript. All authors contributed and approved the manuscript.

## Extended Data

**Extended Data Figure 1.**
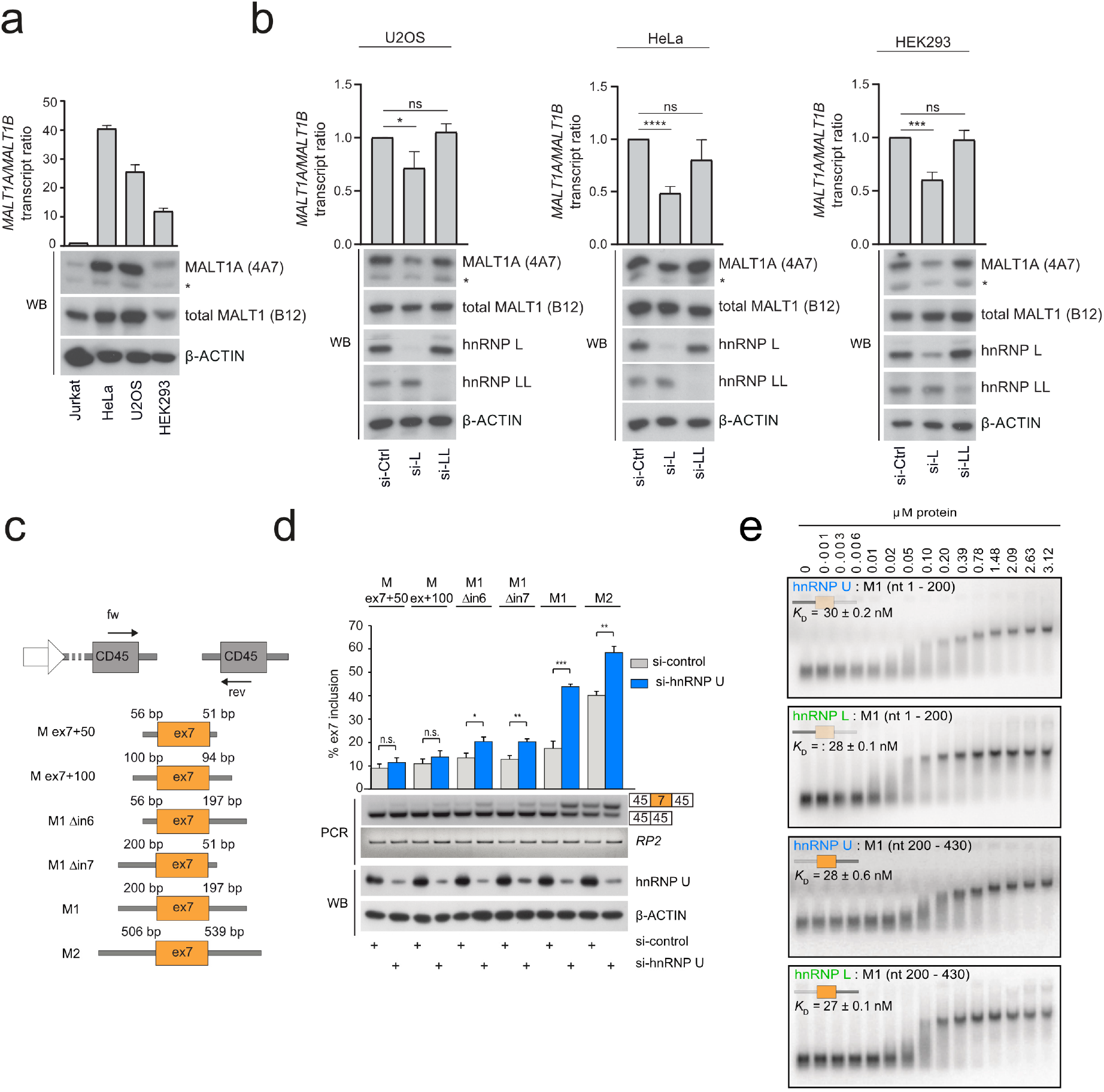
Mapping *MALT1* regions regulated by hnRNP U and L. **a,b**, Quantification of endogenous *MALT1* transcript and MALT1A protein levels upon knockdown of hnRNP L and LL in different cell lines. Asterisk indicates an unspecific band. The antibody used is in parenthesis. **c**, Minigene constructs used. **d**, Minigene splicing assays to evaluate which regions are required for splicing recapitulation relative to endogenous *MALT1*. Data are representative for three independent experiments. Depicted is the mean ± s.d. *p<0.05; **p<0.01; ***p<0.001; ****p<0.0001; NS, not significant; unpaired t-test. **e**, EMSAs showing that hnRNP U and hnRNP L bind with low nanomolar affinity to sub-fragments of the M1 *MALT1* minigene pre-mRNA (M1 nt 1-200 or M1 nt 200-430).

**Extended Data Figure 2.**
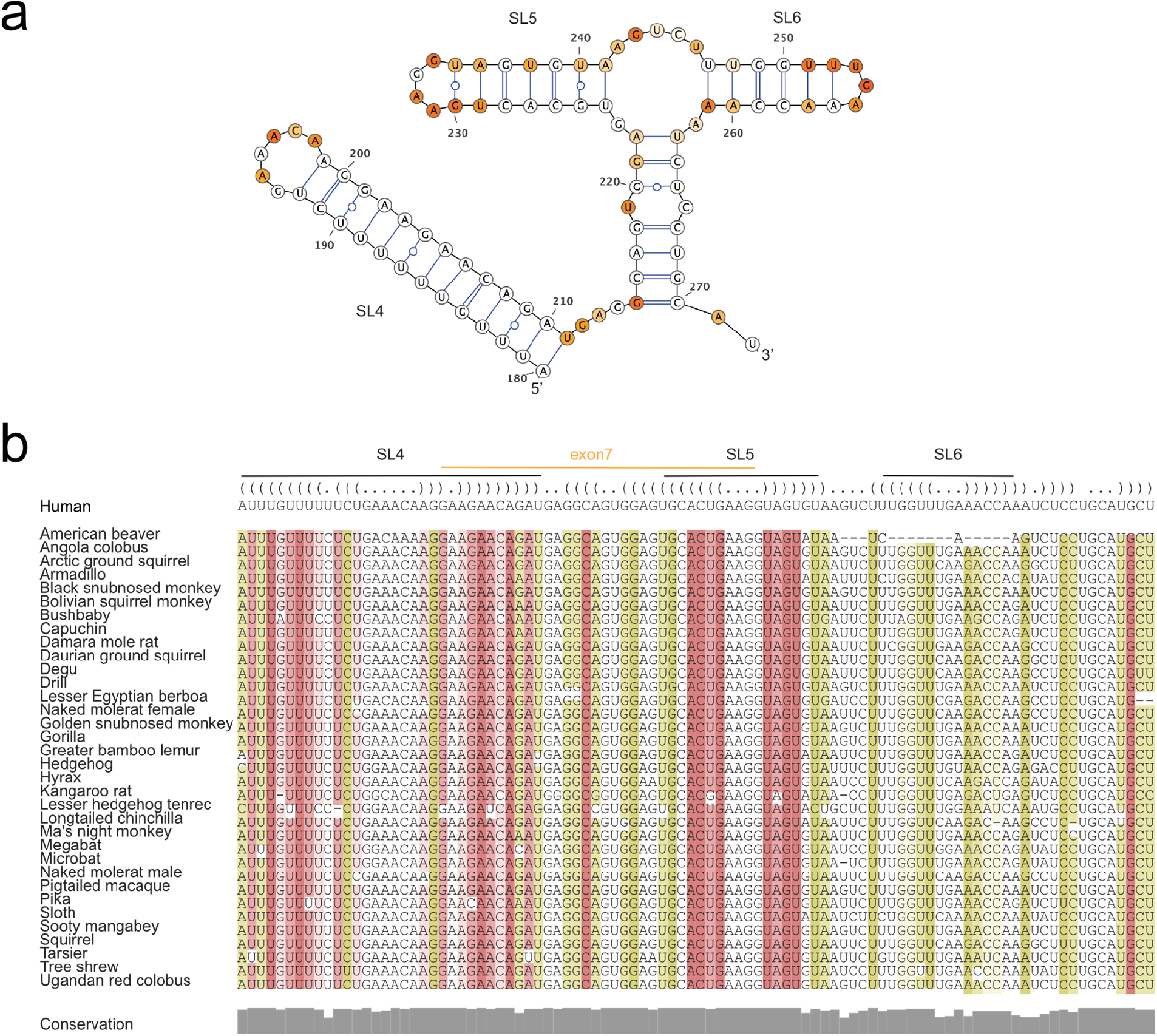
Stem-loop (SL) 4 and the hammerhead, formed by SL5 and SL6, are conserved in mammalian species. **a**, SHAPE reactivities for SL4 and the hammerhead, comprised of SL5 and SL6. Non-reactive, semi-reactive and highly reactive nucleotides are colored white, orange, and red, respectively. **b**, Multiple sequence alignment shows that the RNA sequence and secondary structure for these stem-loops is evolutionary conserved. The alignment is colored using the colorrna.pl of the Vienna package (red marks base pairs with no sequence variation, ochre, green, turquoise, blue, and violet mark base pairs with 2, 3, 4, 5, or 6 different types of base pairs, respectively) ^52^.

**Extended Data Figure 3.**
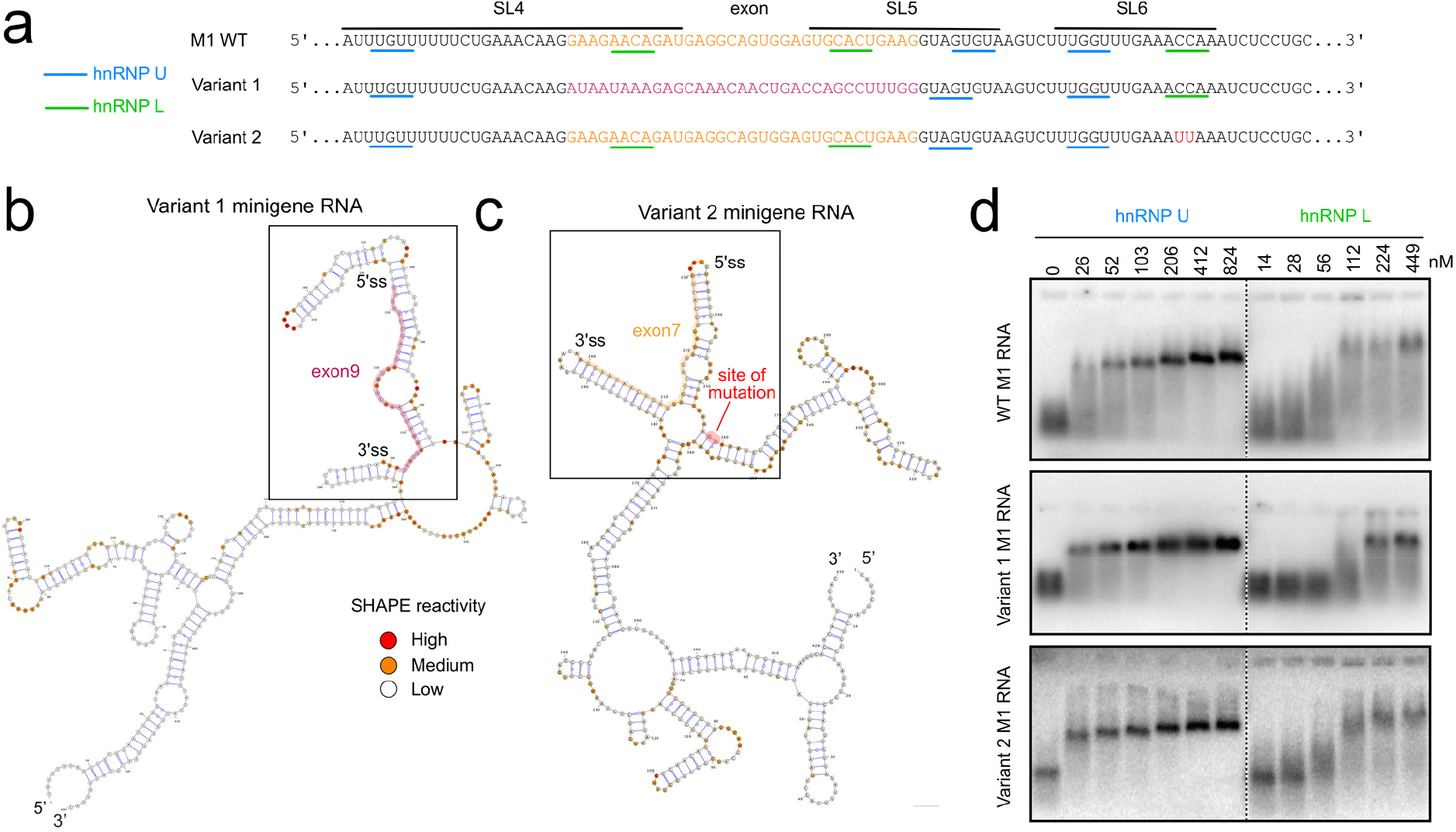
Mutations to the *MALT1* minigene pre-mRNA modify secondary structure, but retain binding with hnRNP U and hnRNP L. **a**, Primary sequences of *MALT1* minigene RNA variants, with mutations colored. **b,c**, Secondary structure of variant 1 and 2 minigene RNAs as determined by SHAPE. **d**, Variants 1 and 2 bind to hnRNP U and hnRNP L comparable to wildtype.

**Extended Data Figure 4.**
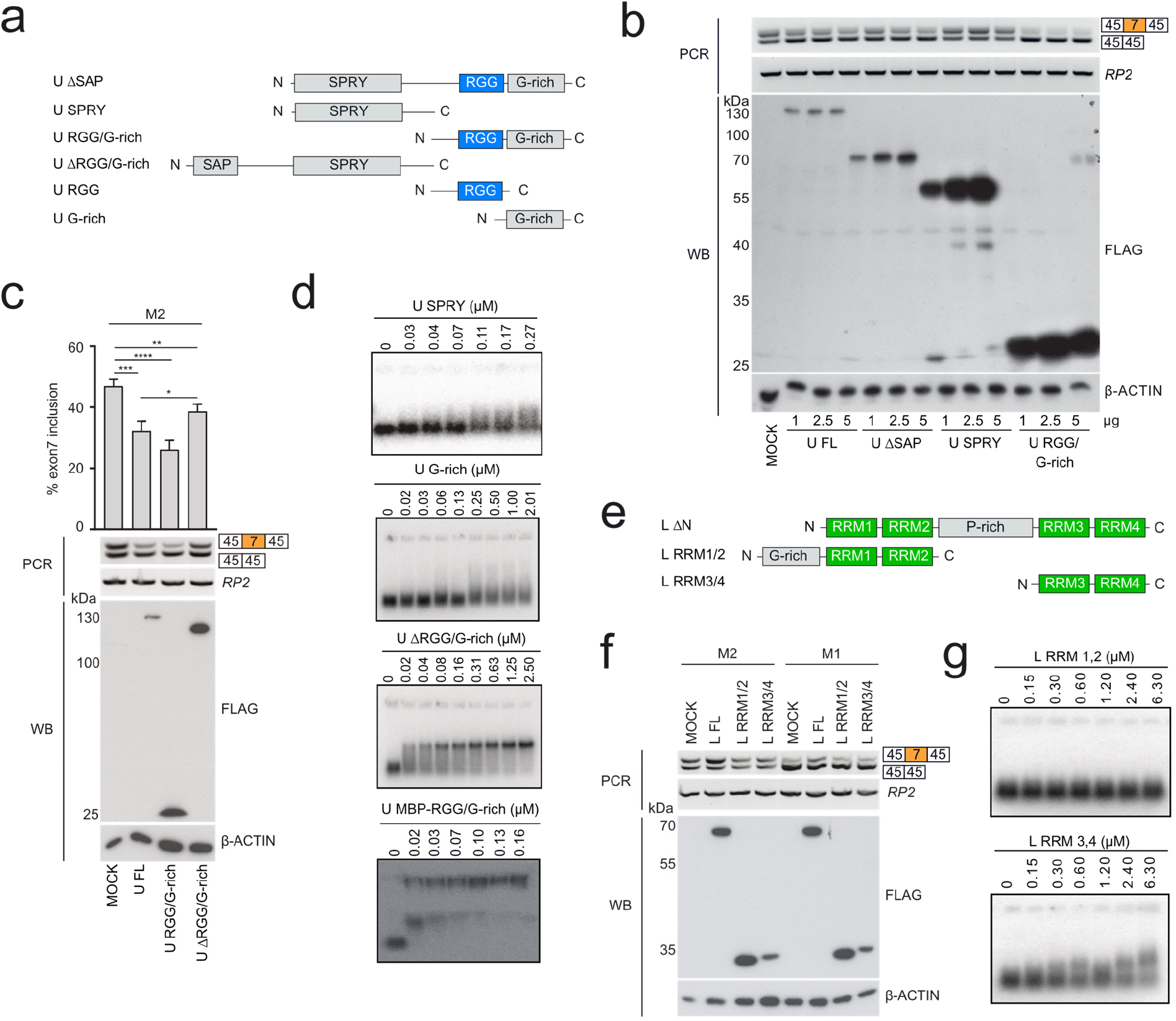
The RGG domain of hnRNP U and the four RRMs of hnRNP L are responsible for binding and regulating splicing of *MALT1* minigene pre-mRNA. **a**, Mapping regions in hnRNP U and L involved in RNA binding. hnRNP U constructs used to identify region responsible for binding RNA. **b,c**, Minigene splicing assays upon overexpression of various hnRNP U constructs. Data are representative for three (**c**) independent experiments. Depicted is the mean ± s.d. *p<0.05; **p<0.01; ***p<0.001; ****p<0.0001; ns, not significant; unpaired Student’s *t*-test. **d**, EMSAs with hnRNP U SPRY, ΔRGG/G-rich, G-rich and RGG/G-rich (with MBP solubility tag) domains with *MALT1* minigene pre-mRNA. **e**, hnRNP L constructs used to identify region responsible for binding RNA. **f**, Minigene splicing assays upon overexpression of various hnRNP L constructs. **g**, Gel shift assays of hnRNP L RRM1,2 and RRM3,4 with *MALT1* minigene pre-mRNA.

**Extended Data Figure 5.**
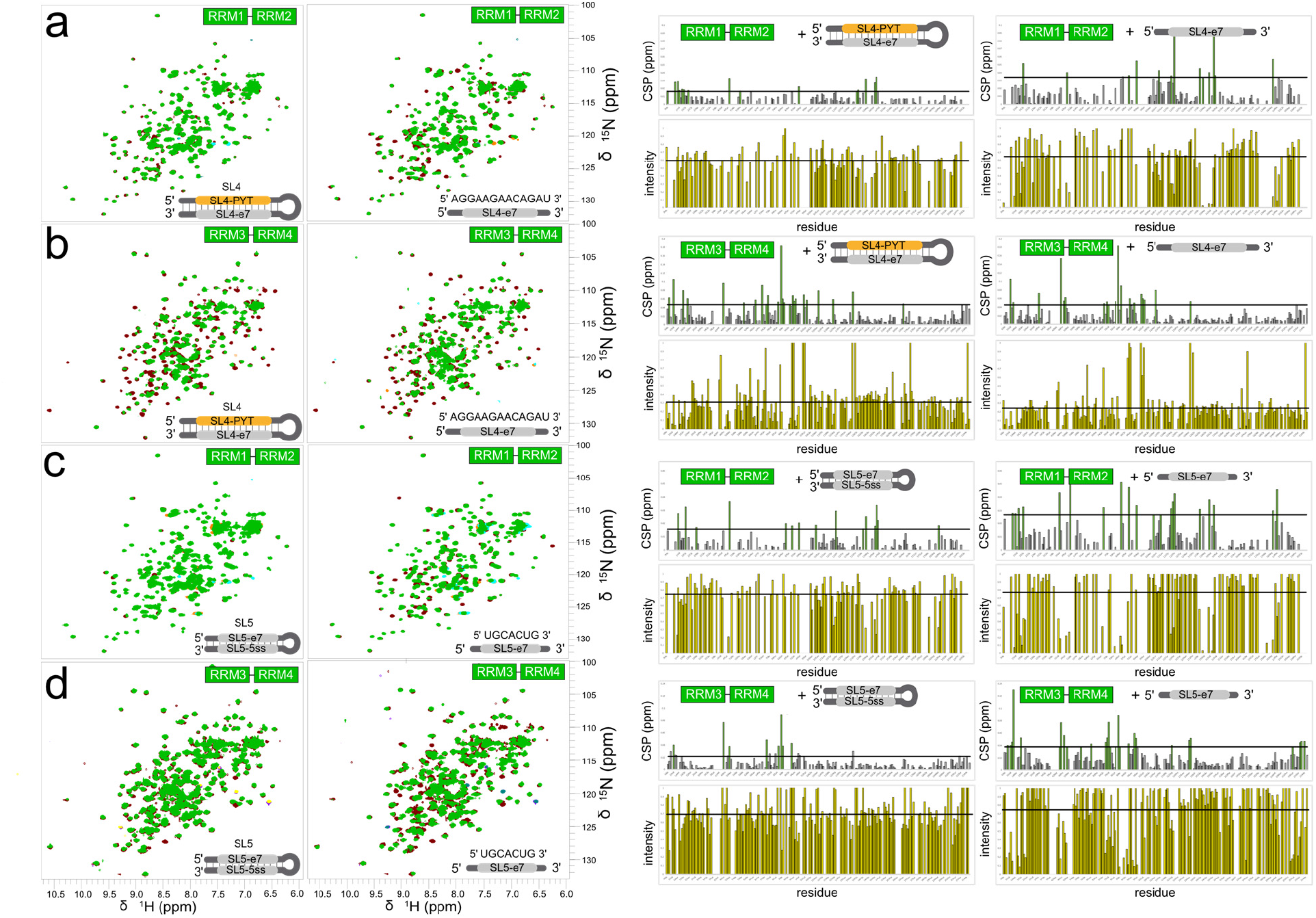
NMR analysis of *MALT1* RNA recognition by hnRNP L. ^1^H,^15^N HSQC spectra of free ^15^N-labeled (**a,c**) RRM1,2 or (**b,d**) RRM3,4 of hnRNP L (maroon) bound to stem-loops or single-stranded components of (**a,b**) SL4 or (**c,d**) SL5 (green) in the *MALT1* pre-mRNA. Chemical shift perturbation (CSP) and intensity plots corresponding to each spectrum are shown on the right. Horizontal lines represent the average CSP plus one standard deviation.

**Extended Data Figure 6.**
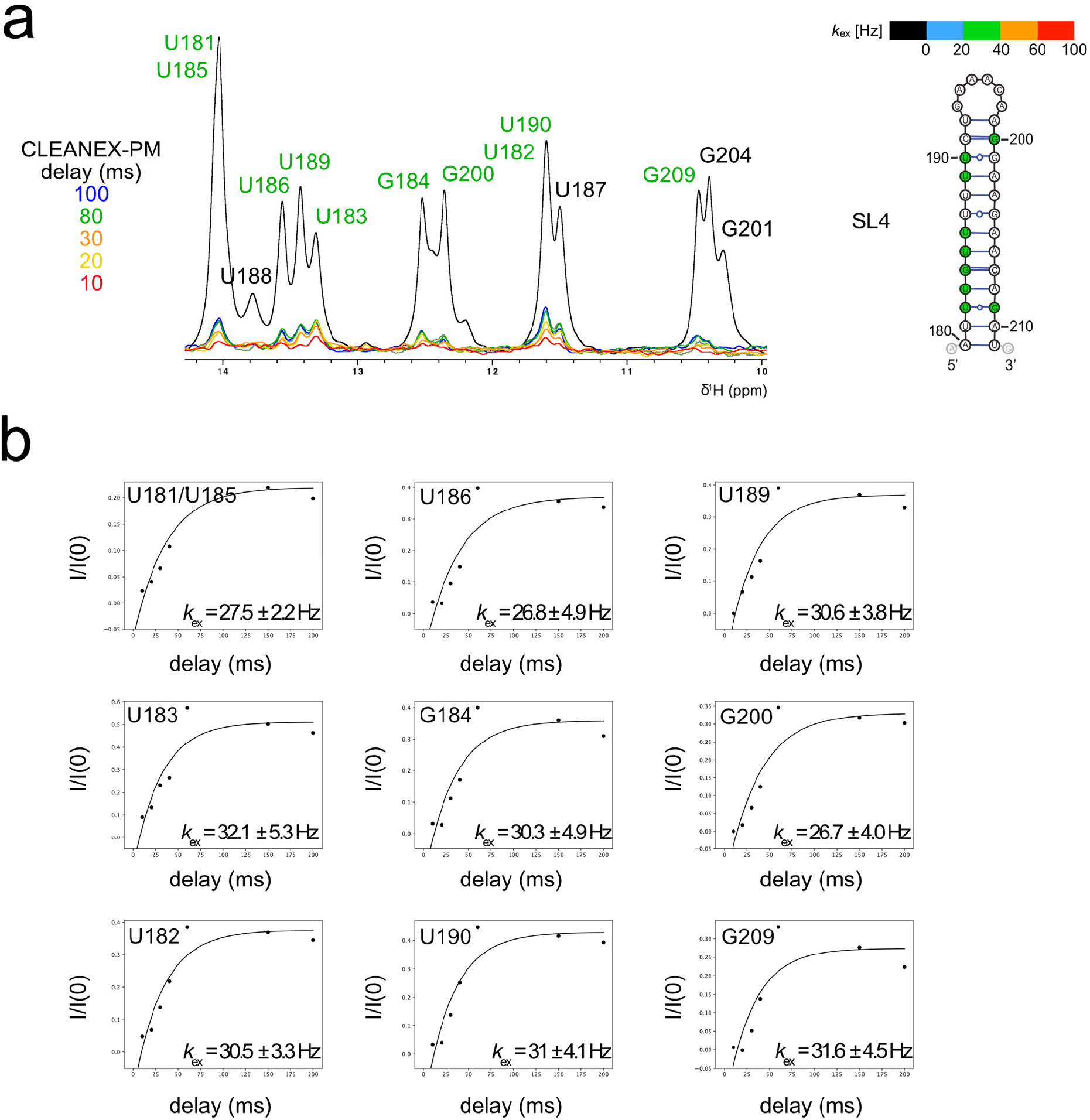
NMR imino proton exchange measurements show reduced stability of *MALT1* SL4 RNA. **a**, Overlay of 1D CLEANEX-PM imino spectra of SL4.Iminos experiencing exchange are labeled in green. **b**, Fitting of peak intensity as a function of CLEANEX-PM mixing time yields water exchange rates *k*_ex_ (in Hz). Error bars indicate the fitting error.

**Extended Data Figure 7.**
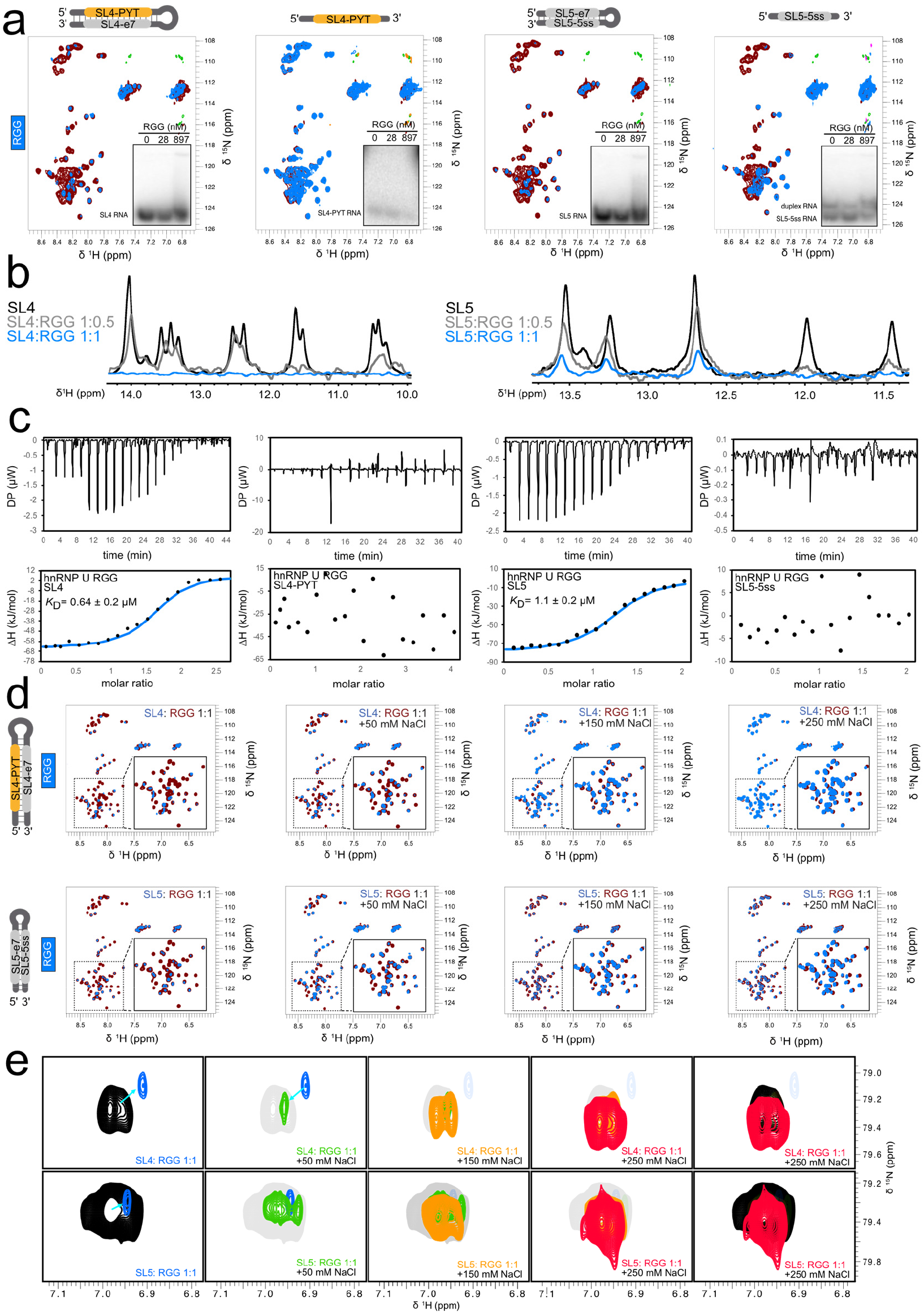
NMR and ITC analysis of *MALT1* RNA recognition by hnRNP U RGG. **a**, ^1^H,^15^N HSQC NMR spectra of the ^15^N-labeled RGG domain of hnRNP U bound to SL4 or SL5 of *MALT1*. **b**, 1D imino titration spectra of SL4 and SL5 (50 µM) in the presence of 0.5 or 1 molar equivalent of the RGG domain of hnRNP U. **c**, ITC heat plot and binding curve of SL4 and SL5 and their single stranded constituents with the hnRNP U RGG domain. **d**, ^1^H ^15^N HSQC spectra of the ^15^N labeled RGG domain of hnRNP U (preformed with either SL4, top, or SL5, bottom) in the presence of increasing concentration of NaCl. **e**, ^1^H ^15^N HISQC spectra of the arginine side chain H^ε^ protons of the hnRNP U RGG domain (preformed with either SL4, top, or SL5, bottom) in the presence of increasing concentration of NaCl.

**Extended Data Figure 8.**
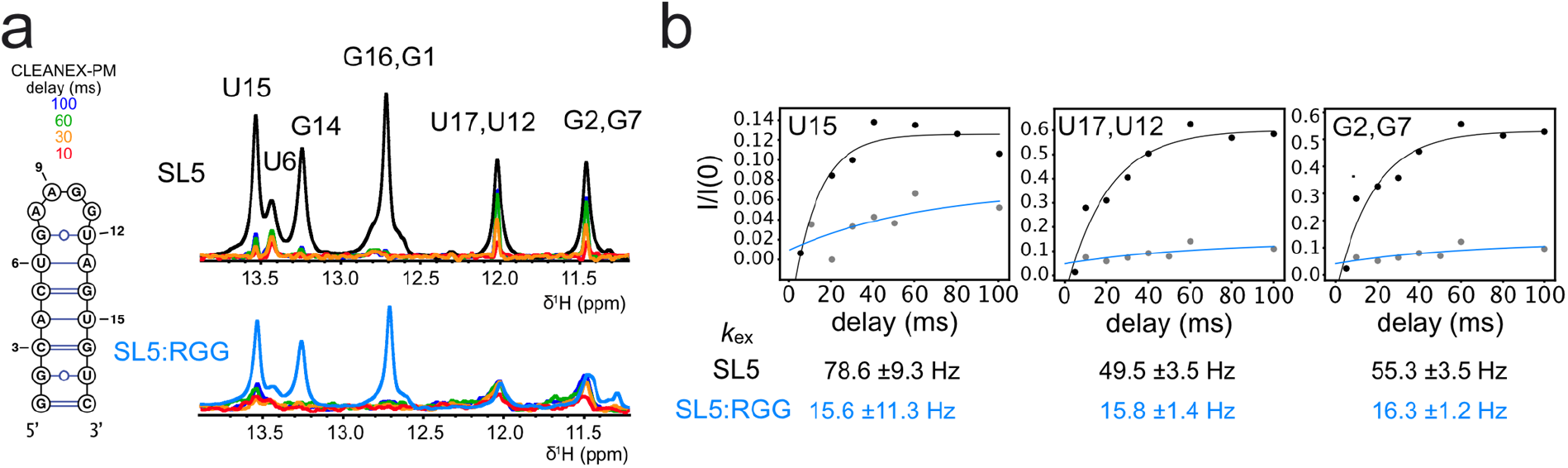
The hnRNP U RGG domain stabilizes secondary structure. **a**, Overlay of 1D CLEANEX-PM imino spectra of SL5 in the absence (black) and presence (blue) of the RGG domain of hnRNP U. **b**, Fitting of peak intensity as a function of CLEANEX mixing time to determined imino proton exchange rates *k*_ex_ (in Hz).

**Extended Data Figure 9.**
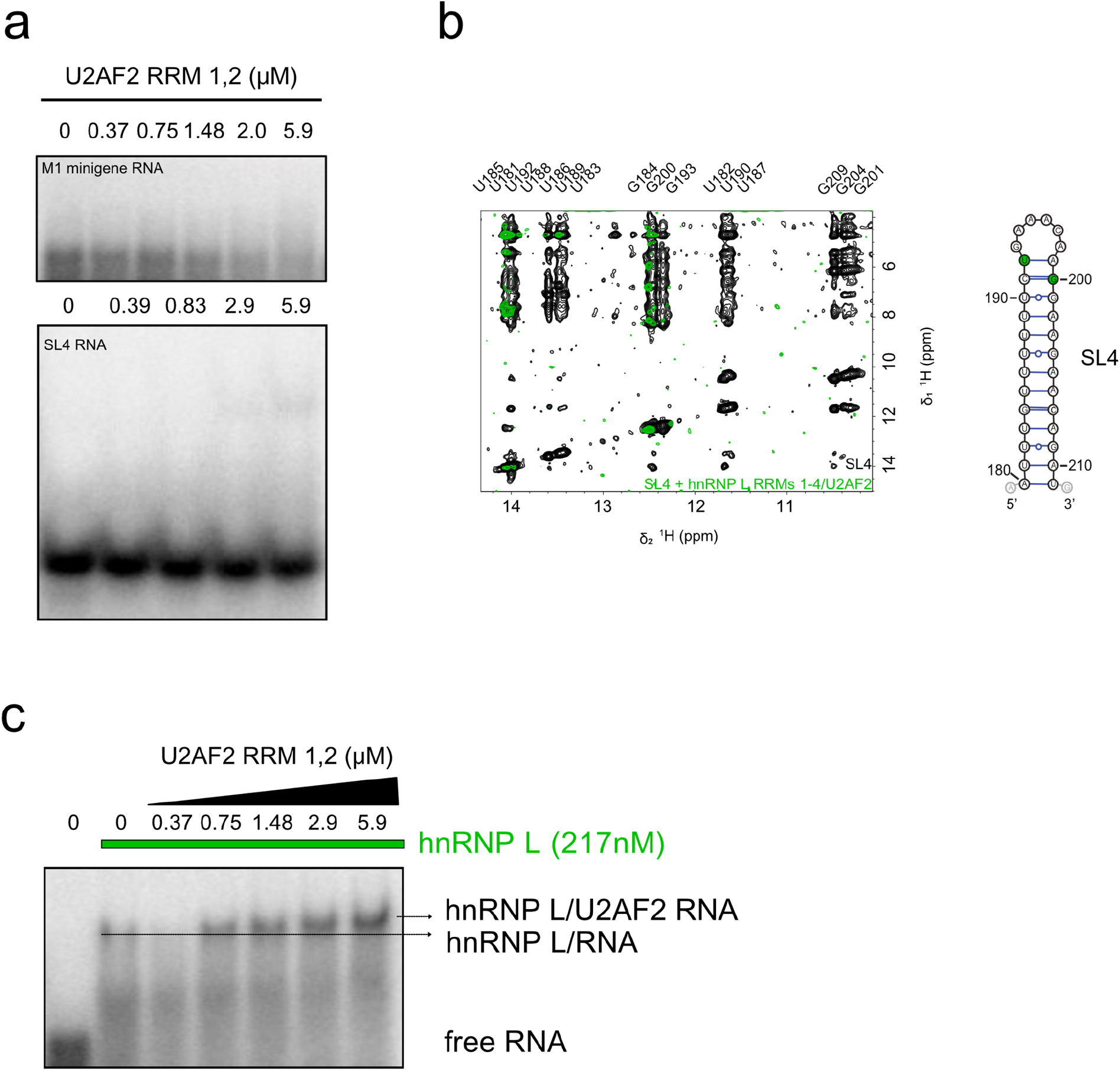
Binding of hnRNP L enables U2AF2 binding to *MALT1* RNA. **a**, EMSA of U2AF2 RRM1,2 with *MALT1* M1 minigene RNA (top) and SL4 (bottom). **b**, Superposition of ^1^H,^1^H NOESY spectra of SL4 RNA in the absence (black) or presence (green) of hnRNP L RRM1-4 and U2AF2 RRM1,2. Imino resonances that remain as part of the protein-RNA complex are colored in green on the secondary structure of SL4. **c**, EMSA of U2AF2 RRM1,2 in the presence of hnRNP L RRM1-4 with M1 minigene RNA.

**Extended Data Figure 10.**
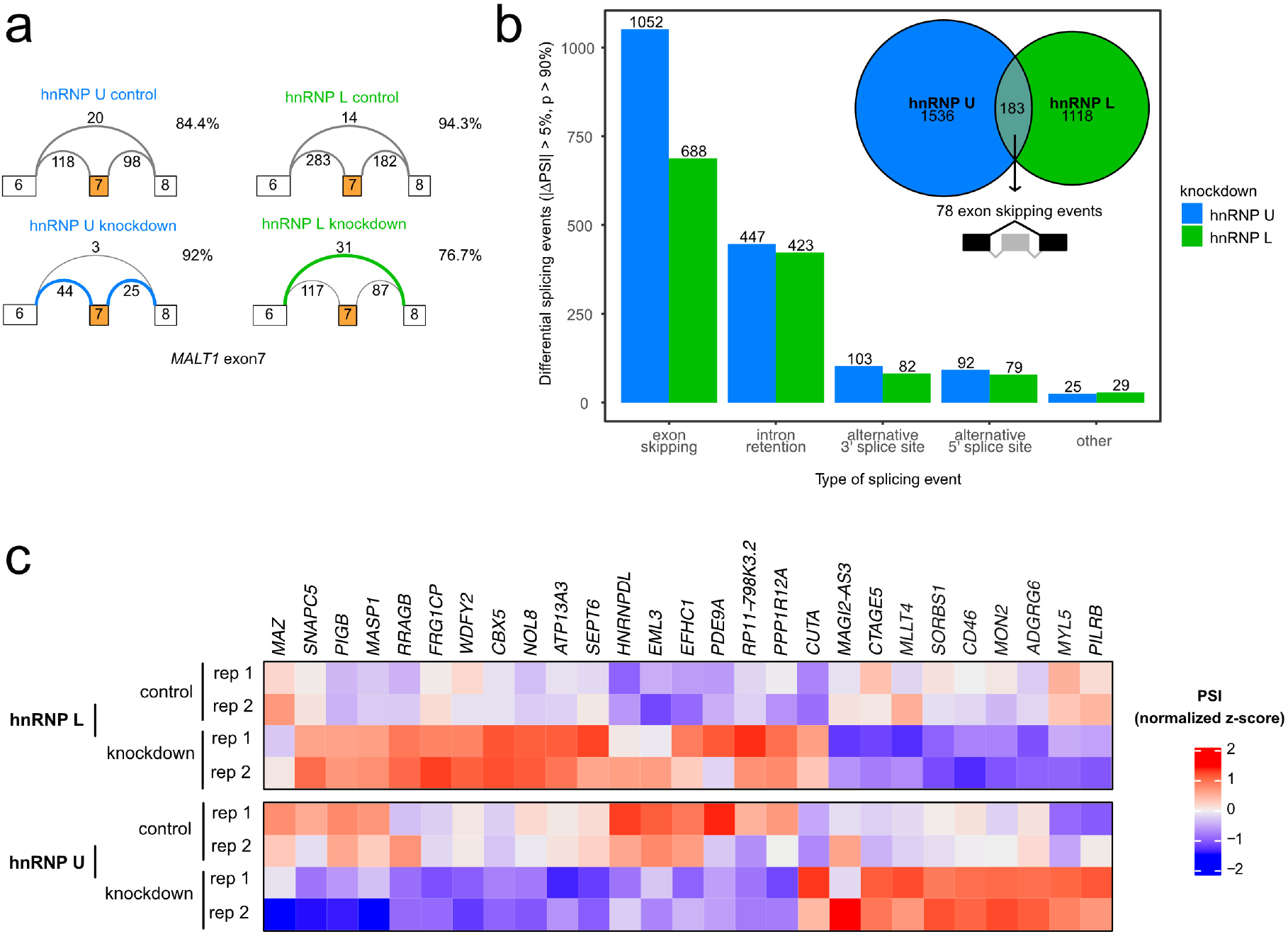
Splicing events regulated by hnRNP U and hnRNP L. **a**, Skipping of exon7 of *MALT1* and its dependence on hnRNP U and hnRNP L is shown from knockdown data available from ENCODE and processed in MAJIQ. Percentage of exon7 inclusion is given for all conditions. **b**, Number of overlapping local splice variations (Venn diagram, number of included exon skipping events given below) and types of splicing events (bar chart) regulated by both hnRNP U and hnRNP L (> 5% change in at least one junction of the local splice variation, 90% confidence interval). **c**, Heatmap showing the comparison of z-score-normalized mean percent selected index (PSI) values for 27 antagonistically regulated cassette exons in hnRNP U and hnRNP L control and knockdown experiments (2 replicates per condition).

**Extended Data Table 1.** Binding affinities for minigene RNAs determined from EMSA.

| RNA construct | hnRNP U |  | hnRNP L |  |
| --- | --- | --- | --- | --- |
| | $K_D^b$ (nM) | $N^a$ | $K_D^b$ (nM) | $N^a$ |
| M1 (nt 1 – 430) | $23.0 \pm 0.5$ | 4.7 | $19.0 \pm 0.2$ | 4.6 |
| M1 (nt 1 – 200) | $30.0 \pm 0.2$ | 2.8 | $28.0 \pm 0.1$ | 2.9 |
| M1 (nt 200 – 430) | $28.0 \pm 0.6$ | 3.1 | $27.0 \pm 0.1$ | 2.8 |

<sup>a</sup> N: Hill coefficient<sup>b</sup> Errors refer to 3 replicates**Extended Data Table 2 | Binding affinities for regulatory RNA elements from ITC**

**Extended Data Table 2.** Binding affinities for regulatory RNA elements from ITC.

| RNA construct | hnRNP U RGG |  |  | U2AF2 RRM1,2 |  |  |
| --- | --- | --- | --- | --- | --- | --- |
| | $K_D^b$ | $\Delta H$ (kJ/mol) | $N^a$ | $K_D$ | $\Delta H$ (kJ/mol) | $N^a$ |
| SL4 | $0.64 \pm 0.2$<br>$\mu M$ | $-76.7 \pm 4.93$ | 1.6 | no binding detected | n.a. | n.a. |
| SL5 | $1.1 \pm 0.2$<br>$\mu M$ | $-85.9 \pm 5.42$ | 1.3 | n.a. | n.a. | n.a. |
| Py-tract | no binding detected | n.a. | n.a. | $2.7 \pm 0.7$ $\mu M$ | $-118 \pm 10.7$ | 1.1 |
| 5' splice site | no binding detected | n.a. | n.a. | n.a. | n.a. | n.a. |
| SL4-ex7 | no binding detected | n.a. | n.a. | n.a. | n.a. | n.a. |
| SL4-5ss | no binding detected | n.a. | n.a. | n.a. | n.a. | n.a. |

<sup>a</sup> N: stoichiometry<sup>b</sup> Errors refer to 2 biological replicates

**Extended Data Table 3.**
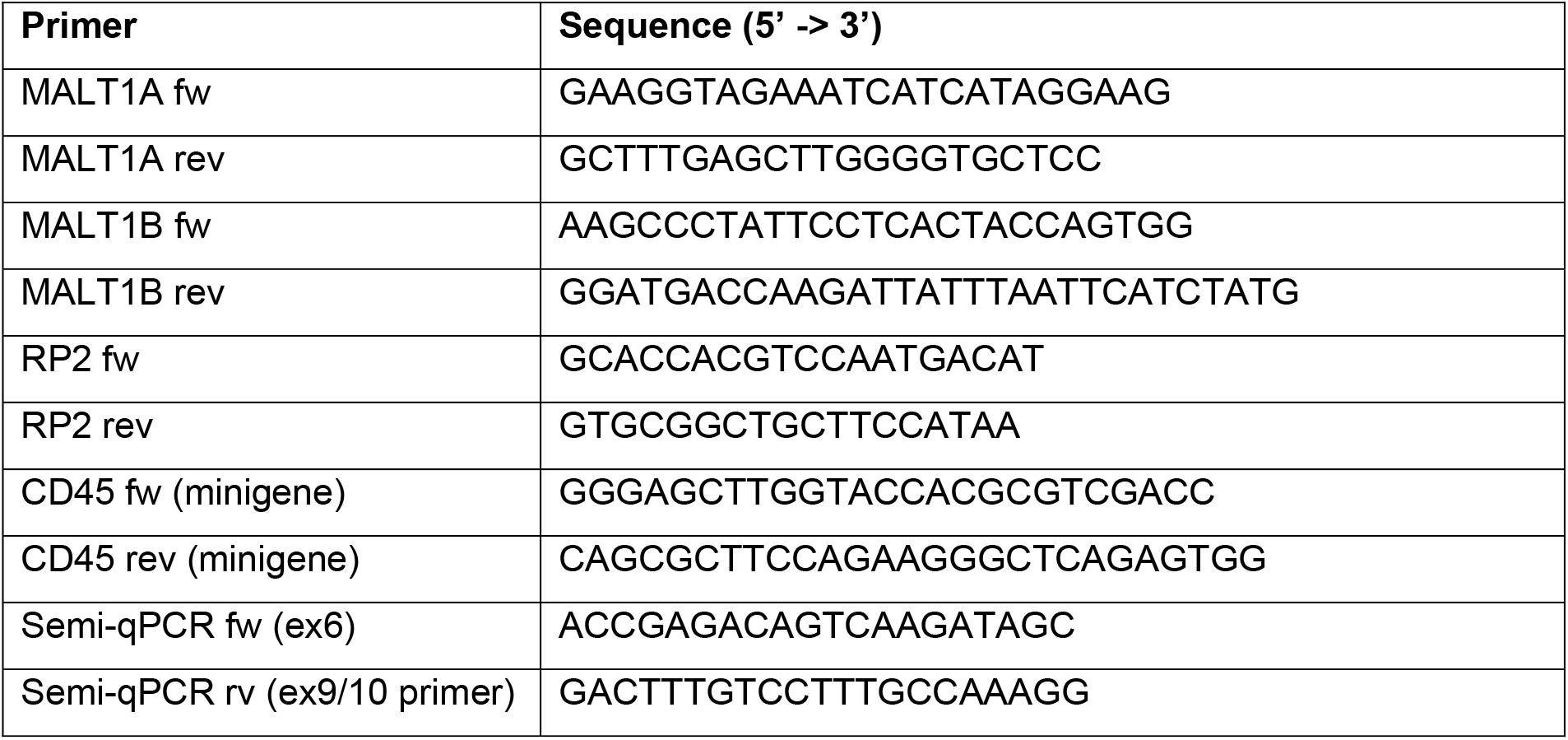
Primers used for qPCR and minigene assays.

**Extended Data Table 4.** Antibodies used for Western blots.

| **Antibody** |
| --- |
| anti-hnRNP U (3G6) Abcam |
| anti-hnRNP L (4D11) Abcam |
| anti-hnRNP LL (4783S) Cell signaling |
| anti-MALT1 (B12) Santa Cruz Biotechnology |
| anti-MALT1A (4A7) HMGU |
| anti-Flag-M2 Sigma-Aldrich |
| anti-β-Actin (C4) Santa Cruz Biotechnology |
| (HRP)-conjugated secondary antibodies (Jackson ImmunoResearch) |

